# Mechanical strain focusing at topological defect sites in regenerating Hydra

**DOI:** 10.1101/2024.06.13.598802

**Authors:** Yonit Maroudas-Sacks, S Suganthan, Liora Garion, Yael Ascoli-Abbina, Ariel Westfried, Noam Dori, Iris Pasvinter, Marko Popović, Kinneret Keren

**Author notes:** Cluster of Excellence, Physics of Life, Technische Universitat Dresden, Arnoldstrasse 18, Dresden, 01307, Germany and Max Planck Institute of Molecular Cell Biology and Genetics, Pfotenhauerstrasse 108, Dresden, 01307, Germany (Yonit Maroudas-Sacks).

## Abstract

The formation of a new head during Hydra regeneration involves the establishment of a head organizer that functions as a signaling center and contains an aster-shaped topological defect in the organization of the supracellular actomyosin fibers. Here we show that the future head region in regenerating tissue fragments undergoes multiple instances of extensive stretching and rupture events from the onset of regeneration. These recurring localized tissue deformations arise due to transient contractions of the supracellular ectodermal actomyosin fibers that focus mechanical strain at defect sites. We further show that stabilization of aster-shaped defects is disrupted by perturbations of the Wnt signaling pathway. We propose a closed-loop feedback mechanism promoting head organizer formation, and develop a biophysical model of regenerating Hydra tissues that incorporates a morphogen source activated by mechanical strain and an alignment interaction directing fibers along morphogen gradients. We suggest that this positive feedback loop leads to mechanical strain focusing at defect sites, enhancing local morphogen production and promoting robust organizer formation.

## INTRODUCTION

Morphogenesis requires elaborate coordination between multiple biochemical and biophysical processes to generate robust, functional outcomes. To achieve this, the system must employ feedback spanning all levels of organization that can relate the emerging patterns with the mechanisms driving their formation (Braun and Keren, 2018; Collinet and Lecuit, 2021). There is a growing appreciation for the role mechanics plays in this process, and in particular the importance of mechanical feedback across scales in directing morphogenetic patterning (Dye et al., 2021; Hannezo and Heisenberg, 2019; Maroudas-Sacks and Keren, 2021). However, despite substantial progress in the field, the mechanisms by which such feedback operate and lead to the emergence of well-defined patterns remain largely elusive.

Hydra is a small fresh-water animal that provides an excellent model system to study the integration between mechanics and other biophysical and biochemical processes in morphogenesis (Braun and Keren, 2018; Gierer, 2012). In particular, Hydra’s capacity for full body regeneration and its short regeneration time facilitate the study of an entire morphogenetic process in a holistic manner. Hydra’s body is composed of a bilayered epithelium that surrounds an internal, fluid-filled lumen. Due to the incompressibility of the fluid, the lumen acts as a hydrostatic skeleton that provides structural support (Kier, 2012). The osmolarity difference between the fresh water environment and the lumen drives an inward osmotic flux (Ferenc et al., 2021; Futterer et al., 2003; Kucken et al., 2008), that inflates the lumen and increases the hydrostatic pressure gradient across the bilayered tissue.

The actomyosin cytoskeleton forms cortical networks and apical junctions in the bilayered epithelium, as well as supracellular actomyosin fibers, known as myonemes, that organize into coherent, parallel arrays spanning the entire Hydra (Aufschnaiter et al., 2017; Livshits et al., 2017). These contractile fiber arrays run parallel to the body axis in the ectoderm, and perpendicular to the axis in the endoderm. All epithelial cells in both layers are excitable, capable of generating calcium-mediated action potentials that activate contraction of the actomyosin myonemes, as in muscles (Agam and Braun, 2023; Braun and Ori, 2019; Campbell et al., 1976; Szymanski and Yuste, 2019). This internal actomyosin-driven force generation together with the hydrostatic pressure, are largely responsible for the deformations and movement of mature Hydra (Szymanski and Yuste, 2019; Wang et al., 2023a) and the shape changes during regeneration (Kucken et al., 2008; Livshits et al., 2017).

A crucial step in Hydra regeneration is the emergence of a head organizer that functions as the main signaling center in mature animals (Bode, 2012). The patterning of regenerating Hydra and the establishment of a new head organizer have mostly been attributed to biochemical morphogens, primarily those associated with the Wnt signaling pathway and its autoregulatory dynamics (Hobmayer et al., 2000; Holstein, 2022; Nakamura et al., 2011). Indeed, local activation of Wnt can promote head organizer formation (Wang et al., 2020) and global Wnt activation results in a multi-headed phenotype (Broun et al., 2005; Ferenc et al., 2021). Despite this progress, axial patterning in Hydra is still not well understood (Holstein, 2022). In particular, even though the significance of the head organizer was recognized over a century ago (Browne, 1909), the mechanisms involved in establishing a new organizer and specifying its location remain obscure (Cazet et al., 2021; Gufler et al., 2018; Suzuki et al., 2023; Tursch et al., 2022).

Regenerating Hydra tissues undergo cycles of osmotic swelling and collapse (Ferenc et al., 2021; Futterer et al., 2003; Kucken et al., 2008). Previous experiments showed that increasing the osmolarity of the external medium leads to diminished osmotic inflations (Futterer et al., 2003; Kucken et al., 2008) and reduced activation of Wnt signaling (Ferenc et al., 2021). In particular, when the osmolarity of the external media and the lumen are comparable, Wnt expression falls off and head regeneration does not occur. While the mechanisms involved are unclear, the repression of the Wnt pathway and the failure to regenerate were attributed to the lack of tissue stretching under isotonic conditions (Ferenc et al., 2021).

Previously we found that sites of topological defects in the nematic order (i.e., in the parallel alignment) of the ectodermal actomyosin fibers emerge early in regenerating Hydra (<24 hours from excision), and coincide with the sites of formation of morphological features (Maroudas-Sacks et al., 2021). Specifically, aster-shaped +1 defects emerge at the future head, and a pair of +½ defects come together in the regenerating foot, in accordance with a memory of body-axis polarity in the parent animal (Shani-Zerbib et al., 2022). Here we study the dynamics of regenerating Hydra spheroids originating from rectangular tissue pieces at high spatiotemporal resolution. We find that tissue deformations are highly non-homogeneous in both space and time, and are strongly-correlated with the nematic organization of the ectodermal actomyosin fibers. In particular, we find recurring tissue stretching concentrated at focus points of the actomyosin fiber pattern. We further find that rupture holes form exclusively at these actin foci, primarily at the future head site. Notably, we show that the actin foci can be identified in the folded spheroid *from the onset of regeneration* and coincide with the location where topological defects emerge and eventually the regenerated head and foot form. The mechanical strain focusing at actin foci is recapitulated by a biophysical model of Hydra tissue mechanics that considers transient activation of contraction in the ectodermal actomyosin fibers.

The colocalization of focused tissue stretching, an aster-shaped +1 defect in the ectodermal fiber organization, and the emergence of a signaling center associated with the new head organizer, suggests a tight coupling between mechanics and signaling. Indeed, we find that disrupting the Wnt signaling pathway using iCRT14 (Cazet et al., 2021; Gufler et al., 2018), or placing tissues in elevated external osmolarity (Ferenc et al., 2021), prevents the formation and/or stabilization of aster-shaped defects and the tissues fail to regenerate. We propose a self-organization mechanism in which actomyosin fiber organization and tissue mechanics are coupled to the dynamics of a biochemical morphogen via a closed-loop feedback mechanism, involving strain-dependent morphogen production at defect sites. We hypothesize that this mechanochemical feedback loop underlies the robust formation and stabilization of the head organizer and use model simulations to demonstrate the viability of this idea, providing the basis for a putative mechanochemical framework for Hydra regeneration.

## RESULTS

### The actomyosin fibers form a reproducible pattern with two foci that become sites of topological defects

The dynamic organization of the ectodermal actomyosin fibers in regenerating Hydra is followed using live imaging of transgenic animals expressing Lifeact-GFP (Methods). Rectangular tissue segments cut from the body of mature Hydra fold and seal into hollow spheroids within a couple of hours (Bode and Bode, 1984; Javois et al., 1988; Livshits et al., 2017) (Fig. 1A,B). This process is highly stereotypical: opposite ends of the tissue stretch to meet, such that the originally head-facing side of the excised tissue comes together with the originally foot-facing side of the tissue (Shani-Zerbib et al., 2022). The excised tissue inherits an array of aligned actomyosin fibers, but during the folding process, the fibers along the edges of the excised tissue disassemble (Livshits et al., 2017; Maroudas-Sacks et al., 2021). The resulting tissue spheroid contains a domain of well-ordered ectodermal fibers corresponding to the more central part of the original excised tissue piece, and a region that lacks supracellular fiber organization (Figs. 1A,B, S1). The ordered domain of parallel fibers spans roughly two-thirds of the tissue circumference in one direction. At either end of this ordered domain (along the fibers’ direction), we find disordered ‘caps’ connected by a disordered ‘bridge’. The two caps are mostly encircled by radially aligned fibers, whereas the connecting bridge is flanked on both sides by fibers aligned parallel to the domain boundary. Overall, the disordered region has a total nematic charge of +2, defined by the fiber organization surrounding it (Fig. 1A). This characteristic pattern allows us to identify two focus points of the actomyosin fiber organization as the centers of the disordered caps (Figs. 1A,B, S1).

**Figure 1.**
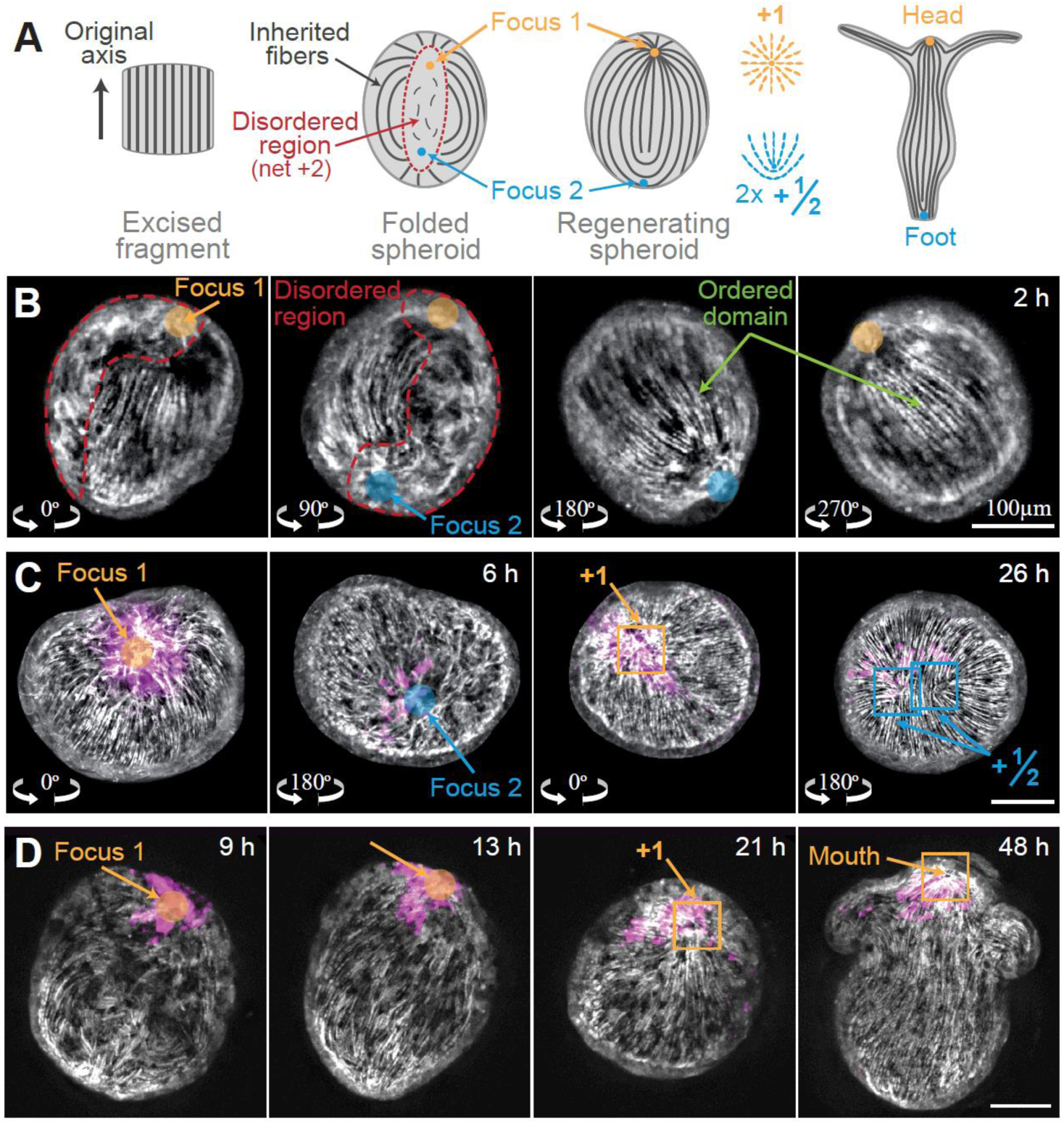
Foci of the actomyosin fiber organization. (A) Schematic of the fiber organization during regeneration. An excised tissue fragment folds into a hollow spheroid that contains an ordered fiber domain and a disordered region. The two actin foci within the initially disordered region develop into an aster-shaped +1 defect at the future head site and a +½ defect pair at the future foot region. (B) Images of a tissue spheroid 2 hours after excision, displaying fiber organization as in (A). The tissue is viewed from 4 directions, rotated 90° from each other, within a square FEP tube (Methods). This characteristic fiber pattern was observed in 46/66 tissue spheroids similarly imaged 2-6 hours after excision. The remaining samples showed consistent behavior, but the pattern was not clear. (C) Images from a time-lapse movie of a regenerating Hydra acquired simultaneously from two opposite sides (Movie 1). The actin organization (gray) is shown with an overlay of the photoactivated dye (magenta; Abberior CAGE 552). Left: the two actin foci within the disordered domain are visible at an early time. Right: the same regions at a later time (as indicated by the tissue label) displaying the characteristic defect configuration at the two foci. (D) Images from a time-lapse movie of a regenerating Hydra showing an actin focus developing into a +1 defect at the tip of the future head. The actin organization (gray) is shown with an overlay of the photoactivated dye (magenta; Abberior CAGE 552), identifying cells in the vicinity of the actin focus at the onset of regeneration as the same cells that later reside at the regenerated head. All images are projected views of the ectodermal basal surface generated from 3D spinning-disk confocal images of regenerating Hydra expressing Lifeact-GFP in the ectoderm.

The early stages of the regeneration process are characterized by an induction of order process, during which the inherited fibers guide the formation of aligned fibers throughout the spheroid, and point topological defects emerge (Maroudas-Sacks et al., 2021). We find that the location of the emerging defects is highly stereotypical, forming at the center of the caps at the two foci of the fiber organization (Figs. 1, S1). Using laser-induced uncaging of a caged dye (Abberior CAGE 552; Methods) (Maroudas-Sacks et al., 2021), we label groups of cells at the cap regions and follow them over time (Fig. 1C,D; Movie 1). The cap closer to the head-facing side of the excised tissue develops a +1 aster-shaped defect at the future head site, whereas in the other cap, a pair of +½ defects form at the future foot region. Meanwhile, the ‘bridge’ between the two caps develops a complete, ordered array of fibers aligned parallel to the inherited fibers along its boundary.

### Mechanical strain focusing at actin foci

The tissue dynamics during regeneration are followed in conjunction with the supracellular actomyosin fiber organization at high spatiotemporal resolution. The Lifeact-GFP probe that labels the actomyosin fibers at the ectodermal basal surface, also binds filamentous actin at cell-cell junctions at the apical surface (Aufschnaiter et al., 2017). We developed an image processing pipeline to separate the fluorescent signal from the apical and basal sides of the ectoderm, allowing us to identify individual cells at the apical surface, while simultaneously following the actomyosin fiber organization (Methods). The regenerating tissue is extremely dynamic, with continuous movement and deformations. The most pronounced deformations are seen when the tissue experiences large-scale, coordinated contractions, that are reflected in extensive, yet transient, distortion of the tissue, primarily along the direction of the ectodermal actomyosin fibers (Fig. 2). During these events, we observe stretching of cells at the center of the disordered cap regions (Fig. 2A), and at +1 defect sites following the induction of order at later stages of regeneration (Fig. 2B). Occasionally we observe ‘doming’, where the stretched tissue exhibits an abrupt curvature change between the core stretched region and the rest of the tissue (Fig. S2), suggestive of inhomogeneous or non-linear material properties (Latorre et al., 2018).

**Figure 2.**
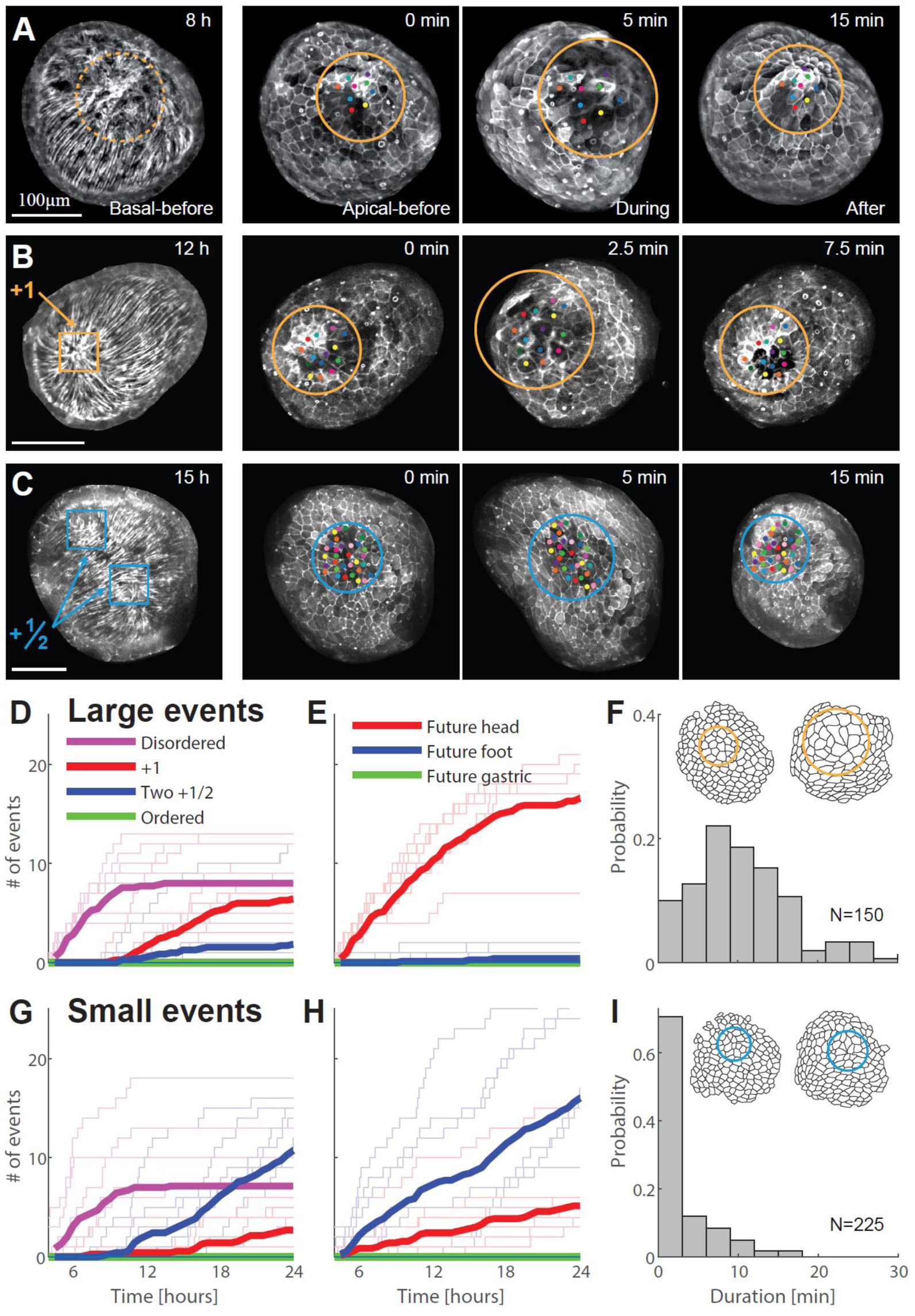
Recurring tissue stretching events at actin foci. (A-C) Images from time-lapse movies showing transient, local stretching events at actin foci: (A) at the center of one of the caps in the early disordered region; (B) at a +1 defect; and (C) at a +½ defect pair. For each event, the left image depicts the ectodermal basal surface, and the right images show the apical surface throughout the event, with colored dots marking manually-tracked cells. All images are projected spinning-disk confocal images of regenerating tissues expressing Lifeact-GFP in the ectoderm. (D-I) Analysis of the distribution of larger (D-F) and smaller (G-I) stretching events. Widespread stretching events are manually classified as large (>∼2-fold cell area change or rupture) or small (<∼2-fold cell area change; Methods). (D) Plot showing the cumulative number of large stretching events observed 4-24 hours after excision, as a function of the fiber organization at the event site. The thin lines depict results from 7 high-resolution movies of regenerating fragments imaged using an up-and-under microscope (Methods), and the thick lines denote the mean value. All samples regenerated, with an average regeneration time of 29±6 hours (mean±std). Local stretching events continue throughout the regeneration process, but become harder to visualize after >20 hours, since the regenerating tissues typically elongate and orient parallel to the imaging plane. (E) Plot showing the cumulative number of large stretching events for the same movies as in D, as a function of the morphological fate of the event site in the regenerated animal. (F) Histogram showing the duration of large stretching events in the first 24 hours after excision. Inset: Schematic showing cell outlines during a large stretching event. (G-I) Plots as in D-F for small stretching events.

To quantify the frequency and localization of stretching events, we use spinning-disk confocal microscopy in a custom up-and-under setup, which enables simultaneous imaging of multiple samples from two opposite sides (Methods). We image at 2-2.5 minutes’ time resolution, which is shorter than the typical event duration (7 ± 6 min, mean ± std, N=375 stretching events; Fig. 2F,I), and manually record tissue stretching events during the first 24 hours following excision (Fig. 2D-I). Large stretching events, defined as incidents in which stretched cells reach >2-fold increase in apical cell area, are observed only at actin foci, mostly at the disordered cap regions early on or later at sites of +1 defects (Fig. 2A,B,D; Movie 2). These large stretching events occur at the future head site, at a rate of ∼1 event/hour (Fig. 2E). Notably, we never observe stretching events in ordered regions, where the fibers are organized in a parallel array, and observe only smaller stretching events (< 2-fold increase in apical cell area) at regions containing +½ defect pairs (Fig. 2C,G). As the regenerating tissue becomes elongated, it often aligns parallel to the imaging plane making it harder to visualize the defect regions, yet pronounced tissue deformations continue. Similarly, in mature animals, contraction of the ectodermal fibers leads to stretching at the +1 defect region and mouth opening (Carter et al., 2016).

The tissue stretching events typically occur simultaneously at both foci, with the future head region consistently exhibiting larger amplitude events compared to the future foot region (Figs. S3, 2E,H). This is true even at early stages of the regeneration process, where both cap regions lack ordered fibers and we are unable to distinguish between them based on the fiber organization. We have previously shown that the future head and foot form in accordance with a memory of body-axis polarity (Javois et al., 1988; Shani-Zerbib et al., 2022). Thus, while the details of how body-axis polarity memory is encoded in the tissue remain unknown, our results indicate that there is a mechanical manifestation of this memory in the tissue deformations, which is apparent already at the earliest stages of the regeneration process (Fig. 2E,H).

The patterns of tissue deformations at the future head and foot regions become more distinct as the fibers organize and form the characteristic defect configuration, with a +1 defect at the future head and a +½ defect pair at the future foot (Figs. 2B,C, S3). Events at the future head region, remain radially symmetric around the +1 defect and are similar in amplitude to earlier events (Fig. 2A,B). However, events localized between two +½ defects exhibit more moderate cell area stretching (Fig. 2C). These difference between the pattern and extent of stretching, can be attributed to the different contractile fiber organization, and particularly the different defect configuration (+1 vs +½ pair, respectively) at the future head and foot regions.

### Characterization of the deformation pattern during stretching events

To quantify the pattern of ectodermal tissue deformations during stretching events, we segment individual cells based on the Lifeact-GFP signal at the apical cell-cell junctions, correcting for the geometrical distortions due to the curved tissue surface (Fig. 3; Methods). As in other epithelial tissues, cell segmentation provides a local measure of tissue deformations (e.g. (Blanchard et al., 2009; Etournay et al., 2015; Guirao et al., 2015; Hashimoto et al., 2015; Priya et al., 2020)). Tracking individual cells in regenerating Hydra is difficult due to the highly dynamic nature of the tissue. However, the actomyosin fiber pattern, which remains stable during stretching events (∼several minutes; Fig. 2F,I), provides a useful frame of reference. We thus use the actin foci as reference points, and quantify the pattern of cellular deformations as a function of distance from these points. Since geometric distances depend on the varying tissue strain, we use graph distance (i.e. the degree of minimal neighbor separation between cells) to quantify the distance from the actin foci (Methods). We measure cell area strain using logarithmic strain, defined as the natural logarithm of the ratio of cell area during the peak of a stretching event with the cell area just before the event started.

**Figure 3.**
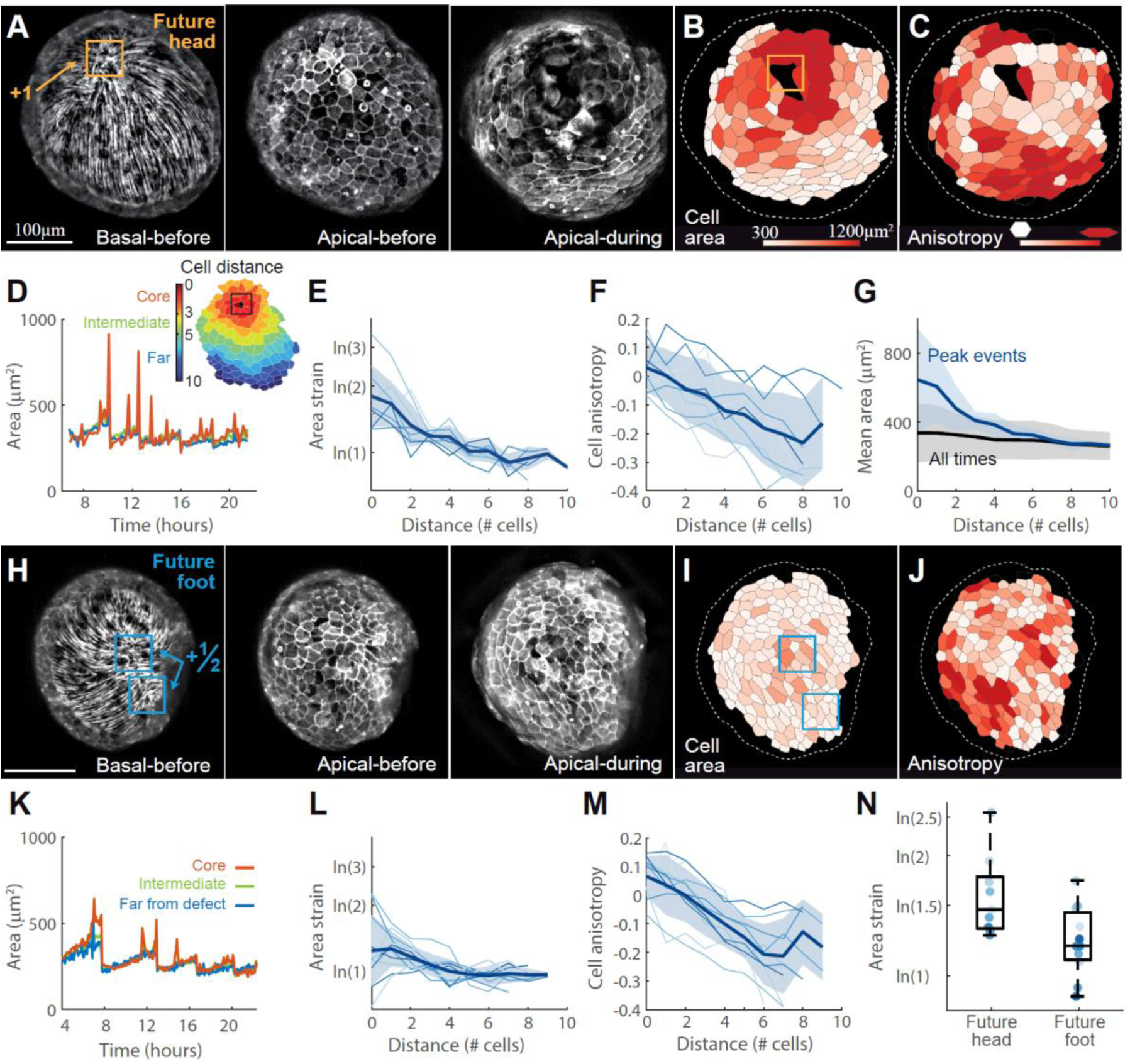
Characterization of mechanical strain focusing at actin foci. (A) Images showing the fiber organization at the ectodermal basal surface (left), and the cellular organization at the apical surface of a regenerating Hydra before (center) and at the peak (right) of a large stretching event at the future head region. (B,C) Cell segmentation maps at the peak of the stretching event shown in A, depicting cells labelled according to their area (B) or cell anisotropy (C; range: 0.1-0.65). (D) Plot of the mean cell area as a function of time, extracted from a time-lapse movie of a regenerating Hydra imaged from the future head side (Movie 3). The observed tissue is divided into three regions: (i) focus region (≤3 cells from focus; red), (ii) intermediate (4-5 cells from focus; green), and (iii) far from the focus (≥6 cells from focus; blue). The focus is defined as the center of the cap region early on, or as the location of the +1 defect once formed. The focus region shows large, transient increases in cell area, which are not seen elsewhere. (E) The logarithmic area strain at the peak of individual stretching events at the future head site (from Movie 3; the color becomes darker with time), as a function of distance from the focus. The bold line represents the mean over all events, and the shaded region represents the standard deviation. (F) Radial cell shape anisotropy as a function of distance from the focus, measured as the magnitude of the cell anisotropy tensor along the vector pointing towards the focus (Methods). Events, mean and standard deviation are represented as in E. (G) Mean cell area as a function of distance from the focus. The cell area is averaged over frames at the peak of stretching events (blue), or over the entire movie (black; mean±std). (H-M) Images and graphs showing the tissue dynamics in a regenerating Hydra spheroid at the future foot site (Movie 4) in analogy to (A-F). The focus is defined as the center of the cap region early on, or as the mid-point between the location of the two +½ defects once formed. (N) Bar plot comparing the logarithmic area strain at the focus regions for all events recorded in Movies 3 and 4 (N=9 and N=11 events at the future head and foot sites, respectively). The mean and standard deviation are shown together with the results from individual events (dots).

We find that area strain during stretching events is highest at the actin focus at the future head region, located early on at the center of the cap region in the disordered domain and later at the +1 defect site (Fig. 3A,B). The area strain decays within a distance of ∼3-5 cells from the foci (Fig. 3D,E). The strain amplitude at the core varies between events, with an average of ∼ln(2), i.e. a two-fold increase in cell area (Fig. 3E). At the same time, cells away from the focus deform anisotropically, contracting along the direction of the ectodermal fibers (Fig. 3C,F). We quantify the degree of cell anisotropy using the cell shape tensor **Q** (Merkel et al., 2017) (Methods). The deformation pattern at the future foot side that contains a +½ defect pair exhibits only moderate local area stretching at the core (Fig. 3H-N). The cells in the region between the two defects, transiently contract along the fibers’ direction, similar to what is observed in the ordered fiber arrays in the future gastric region.

The observed tissue stretching is not only localized in space, but also in time, with rapid stretching events typically separated by considerably longer time intervals (Fig. 3D,K). Notably, the correlation between cell area deformations and fiber organization (Fig. 3E,F,L,M), is not apparent during most of the regeneration process. In particular, the time-averaged changes in cell area strain around defect sites are substantially smaller than the changes observed during stretching events, and within the typical range of variation in cell area (Fig. 3G). Thus, while the area strain experienced by regenerating tissues during stretching events generates a clear mechanical signature at defect sites, these enhanced tissue deformations are transient and are not apparent upon time averaging. This suggests that if these tissue deformations provide relevant mechanical cues at the future head site, the response to these cues must be non-linear, accentuating the large yet transient strains encountered at defect sites.

### Rupture holes in regenerating Hydra spheroids form at actin foci

Regenerating Hydra spheroids exhibit cycles of swelling and collapse driven by osmotic influx of fluid into the lumen followed by tissue rupture and fluid release (Ferenc et al., 2021; Futterer et al., 2003; Kucken et al., 2008; Wang et al., 2019). Tissue ruptures typically involve both layers of the epithelium. This can be shown directly by imaging the ectoderm and endoderm simultaneously (Carter et al., 2016), as well as indirectly by observing fluid efflux accompanied by a reduction in lumen volume (Ferenc et al., 2021; Futterer et al., 2003; Kucken et al., 2008; Maroudas-Sacks et al., 2021; Wang et al., 2019). Tissue ruptures could have various physiological implications including, e.g., release of internal pressure (Stokkermans et al., 2022), changes in the electric potential gradient across the tissue (Macklin and Josephson, 1971), and induction of a wound-healing response in the tissue (Tursch et al., 2022).

Our high-resolution imaging allows us to visualize the formation of rupture holes and characterize their location and the associated tissue deformations. We find that osmotic inflations involve gradual, roughly homogeneous increase in cell area throughout the ectodermal shell (Figs. 4A). However, rupture holes form exclusively at actin foci, and their formation is preceded by focused tissue stretching of the future hole site (Fig. 4; Movies 5,6). The rupture holes recur at the actin foci sites, and never appear in regions with ordered fibers (Fig. 4A,F). This is true from the very first ruptures that occur in the folded spheroid, contrary to what has been reported previously based on lower-resolution imaging (Wang et al., 2019). We further observe that the formation of rupture holes is biased toward the future head region (Fig. 4G), which also exhibits more extensive tissue stretching (Fig. 3N).

**Figure 4.**
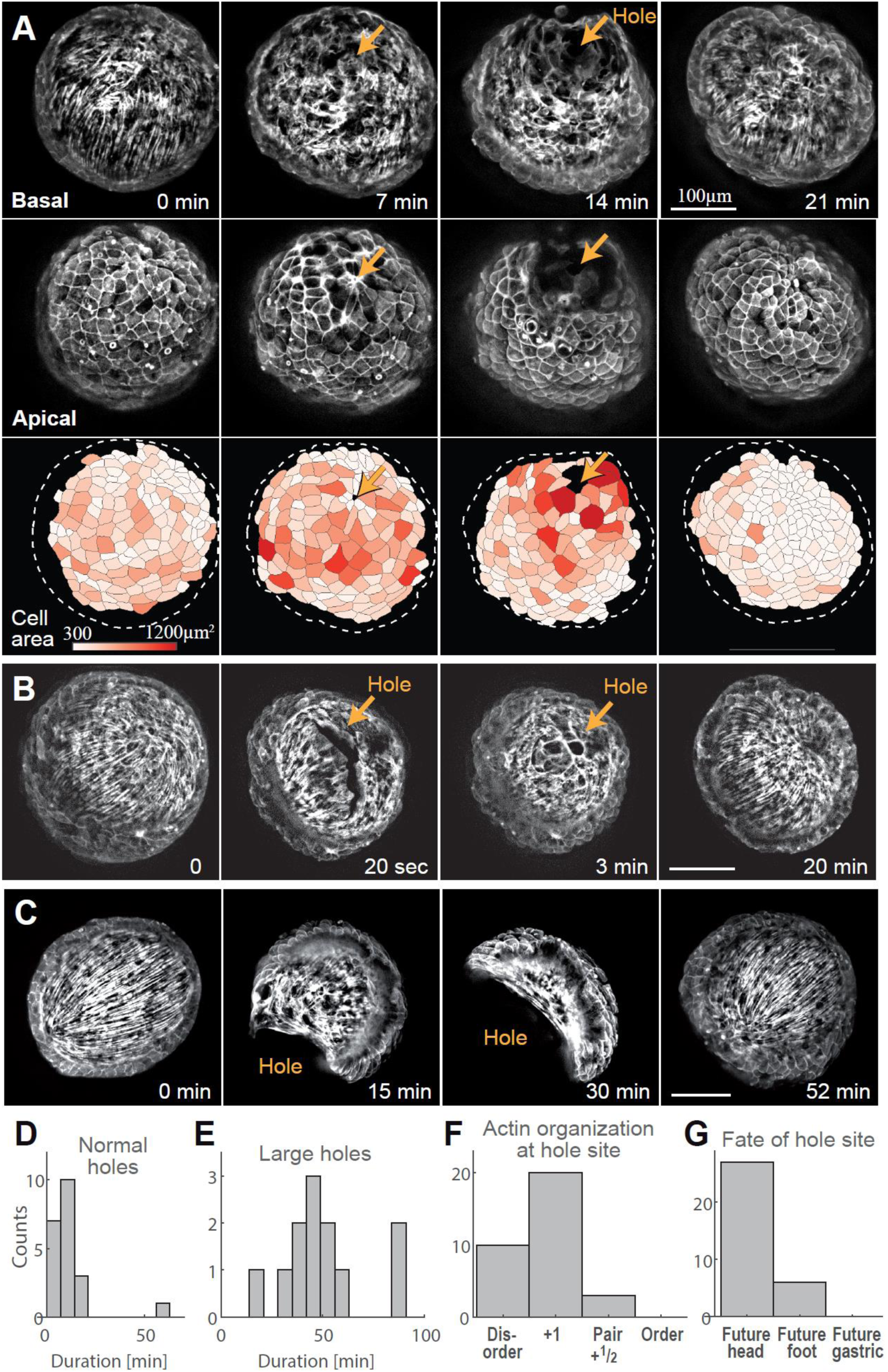
Rupture holes in regenerating tissue spheroids form at actin foci. (A-C) Examples of the opening and resealing of rupture holes at the future head site. The images (B,C and top row in A) show the ectodermal basal fibers. In A, the middle row also shows the ectodermal apical surface, and the bottom row shows segmented cells color-coded according to their area. (B) Images from a fast time-lapse movie showing a rupture that initiates as an elongated crack and rapidly becomes circular (Movie 5). (C) Images from a time-lapse movie showing the opening of an extremely large rupture hole (Movie 6). (D) Histogram of the duration of small rupture holes (as in A,B). (E) Histogram of the duration of large rupture holes (as in C). (F) Bar plot showing the local actin organization at rupture hole sites. (G) Bar plot showing the morphological outcome of rupture hole sites. The data in D-G is based on N=33 rupture holes identified during the first 24 hours after excision in 19 time-lapse movies of regenerating fragments.

The mechanical strain focusing at future rupture sites and compression of cells in the ordered regions far from these sites (Fig. 4A), is analogous to the cellular deformation patterns observed during stretching events (Figs. 2,3). As such, our observations indicate that the tissue ruptures are not the result of random, local failure following homogeneous osmotic inflations as previously thought (Ferenc et al., 2021; Kucken et al., 2008; Wang et al., 2019). Rather, the ruptures form when large-scale actomyosin fiber contractions occur in an osmotically-inflated tissue spheroid. In this case, the contractions further increase the isotropic pressure in the lumen (Stokkermans et al., 2022), and more importantly, generate an additional inhomogeneous in-plane stress that is strongest at the actin foci (see also modeling section below). Rupture hole formation occurs at the site of highest in-plane stress, and hence is localized at the actin foci (Fig. 4F).

Rupture events typically involve disruptions of the epithelial bilayer along cell-cell contacts, generating fractures along cell boundaries (Fig. 4B), as observed in other tissues (Bonfanti et al., 2022). Immediately after their formation, rupture holes can appear as elongated cracks along the junctions connecting adjacent cells (Fig. 4B). These cracks round up within a short time (<1 min) and acquire a smooth boundary, spanning multiple cells, with an enriched actomyosin ring lining the edge of the hole (Movie 5). The rupture holes typically reseal within ∼10min (Fig. 4D). The formation of rupture holes is reminiscent of the mechanism of mouth opening in adult Hydra, which similarly involves tearing the epithelium along cell-cell junctions at a well-defined location that is a focal point of the ectodermal actomyosin fibers (Carter et al., 2016; Wang et al., 2019).

An interesting phenomenon that is observed in about half of the regenerating tissues (8/13 samples observed) is the opening of extremely large rupture holes in the ectoderm, that can even reach the entire spheroid’s circumference (Fig. 4C; Movie 6), and remain open for nearly an hour (Fig. 4E). Remarkably, despite the considerable deformation, the spheroids reseal and appear to recover their pre-rupture organization and eventually regenerate successfully. We suspect that these extremely large ruptures may occur when the ectodermal layer partly detaches from the mesoglea due to the strong shear forces generated by the contraction of the ectodermal fibers. The large opening in the ectoderm in this scenario reflects transient sliding of the ectoderm relative to the endoderm, which subsequently recovers.

### Theoretical modeling of mechanical strain focusing at actin foci

The observed pattern of cellular deformations during large stretching events can be understood intuitively by assuming that the Hydra tissue is elastic (Perros et al., 2023) and that contraction of the ectodermal actomyosin fibers is the primary source of force generation during these events. Upon activation, the aster-shaped +1 defect regions will focus stress (and hence strain) at the defect site, due to contraction of the surrounding radially-oriented fibers. Similar stress focusing is expected at the center of the cap regions, which are encircled by radially-oriented fibers from nearly all directions (apart from the disordered bridge). The fiber organization in the vicinity of +½ defect pairs is expected to be less efficient at focusing stress, and no strain focusing is expected in ordered regions with a parallel fiber array. While the perpendicularly-oriented endodermal fibers also generate forces, we neglect their contribution during stretching events. This assumption is justified by our observations that the tissue rapidly contracts primarily along the direction of the ectodermal fibers during these events (Fig. 3F,M). This is also consistent with recent studies of the behavior of mature Hydra showing that the ectodermal fibers are dominant during large contractions (Wang et al., 2023a).

To corroborate our intuitive understanding of mechanical strain focusing during stretching events, we develop a biophysical model of Hydra tissue mechanics (Fig. 5A; SI). We describe the regenerating Hydra as a deformable shell with an embedded nematic field describing the actomyosin fiber orientation, which encapsulates a fluid-filled lumen. The deformable shell is realized as a collection of cells specified by the positions of their vertices. As such our model is a generalization of 2D vertex models (Farhadifar et al., 2007; Honda and Eguchi, 1980) to curved closed surfaces. The tissue mechanics are described by an energy function *E*, accounting for cell area elasticity, cell-cell adhesion, and cell perimeter elasticity, as in standard vertex models (reviewed in (Alt et al., 2017; Fletcher et al., 2013)). The tissue deformations conserve the volume of the fluid-filled lumen. Furthermore, since we do not explicitly model the tissue thickness, we include a bending energy term that penalizes tissue curvature. We describe the embedded nematic field by assigning a two-dimensional tensor to each cell in the cell-tangent plane. Finally, to emulate stresses generated by fiber contraction, we introduce active cellular stresses, oriented along the nematic field, with a tunable magnitude *ζ* (Comelles et al., 2021). The model parameters are chosen to reproduce the elastic behavior of regenerating Hydra and the magnitude of deformations observed experimentally (SI).

**Figure 5.**
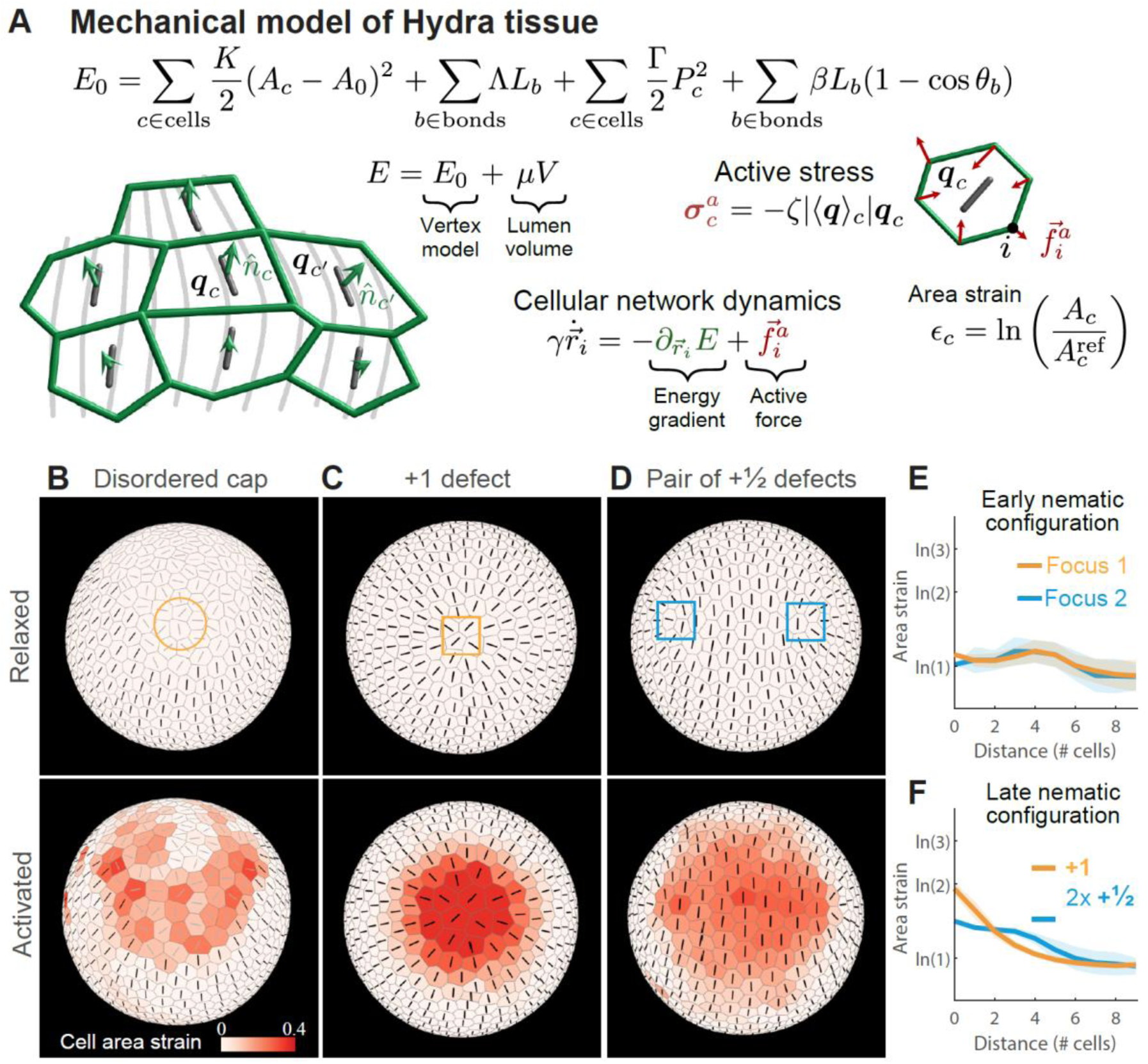
Model of Hydra tissue mechanics recapitulates strain focusing at actin foci. (A) Hydra tissues are described as deformable elastic shells of cells using a generalization of the vertex model. The cellular network geometry is specified by the positions of cell vertices (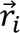) and its dynamics are driven by gradients of an energy function *E* and active stresses generated by the embedded nematic field ***q***_*c*_. Motion of the cellular network is constrained by the lumen volume *V* through a Lagrange multiplier (*μ*). *A*_*c*_ and *A*_0_ are the cell areas and preferred cell area, *K* is the area elastic modulus, *Λ* and *L*_*b*_ are the bond tension and bond length, *Γ* and *P*_*c*_ are the perimeter elastic modulus and cell perimeter, *β* is the cell-cell bending modulus and *θ*_*b*_ is the angle between normal vectors of neighboring cells that share bond *b* (Methods). (B) Mechanical strain focusing at one of the cap regions in a spheroid with a large disordered domain, emulating the fiber organization in regenerating fragments at early stages of the regeneration process. (C,D) Mechanical strain focusing in a spheroid with an ordered nematic array, emulating the fiber organization in regenerating fragments at later stages of the regeneration process. The strain patterns around the +1 defect on one side (C) and around the +½ defect pair at the opposite end (D) are shown. In (B-D), the top panels depict the relaxed configurations before the events, and the bottom panels show the same regions during the peak of the stretching events, with cells color-coded according to the logarithmic cell area strain and the orientation of the nematic depicted. (E,F) Graphs of the simulated logarithmic cell area strain during the peak of a stretching event as a function of distance from the focus. Results are shown for regenerating spheroids at early times around the two foci in the cap regions (E), and at later times around the +1 defect and +½ defect pair (F). The cell area strain is defined as 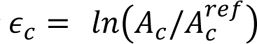, where *A*_*c*_ is the cell area at the peak of the event and 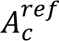 is the initial cell area.

We use this model to study strain patterns induced by individual tissue stretching events (Fig. 5, Movies 7,8). Starting from an initially relaxed state we emulate global contraction activation by introducing a pulse of active stress with magnitude *ζ*_*M*_ (Fig. S4; SI), and duration Δ*T*_*ζ*_. Since the time scale for stretching events (minutes; Fig 2F,I) is shorter than the time scale for fiber reorganization (hours) (Maroudas-Sacks et al., 2021), we assume that the nematic field describing the fiber orientation does not evolve during these events (apart from being conveyed by the cells as the tissue deforms). We consider two particular configurations of the nematic field that recapitulate the observed fiber patterns during early and later stages of regeneration, respectively. For early stretching events, we consider a spheroid with a large disordered domain of net charge +2 surrounded by a fully ordered region (Fig. 5B). For later events, following the induction of order, we consider a spheroid with fully-ordered fibers except for a +1 defect on one end and a +½ defect pair at the opposite end (Fig. 5C).

Our simulations recapitulate the observed deformation patterns, with cell stretching concentrated at the two actin foci, and cell compression along the fibers’ direction in the ordered regions between the foci (Fig. 5B-D, Movies 7,8). Since the fiber organization at early stages of the regeneration is similar around the two foci, the simulated mechanical strain focusing has comparable magnitudes in both foci (Fig. 5E). At later stages of regeneration, when the nematic pattern becomes asymmetric, with a +1 defect at the future head region and a +½ defect pair at the future foot, the model accounts for the asymmetric deformation pattern observed, exhibiting pronounced stretching at the +1 defect site (Fig. 5C,F, Movie 8), and more moderate stretching at the +½ defect pair (Fig. 5D). Overall, these results demonstrate that considering regenerating Hydra as elastic shells with an incompressible lumen that experience global fiber contraction, is sufficient to account for the tissue deformations during stretching events.

### Inhibition of the Wnt pathway disrupts the formation of aster-shaped +1 defects

The coincidence of a +1 defect with the signaling center at the head organizer in mature Hydra (Fig. 6A), suggests an intimate coupling between mechanics and biochemical signaling. To explore this interplay, it is useful to perturb the Wnt signaling pathway, which is considered to be the main activator associated with organizer formation, and characterize the actin fiber organization, tissue dynamics and regeneration outcome. Previous work has shown that local upregulation of Wnt leads to rearrangements of the ectodermal actomyosin fibers toward regions of high Wnt and the formation of aster-shaped +1 defects, both during budding (Aufschnaiter et al., 2017), and following grafting of an excised organizer or a piece of tissue overexpressing Wnt onto a host tissue (Wang et al., 2020).

**Figure 6.**
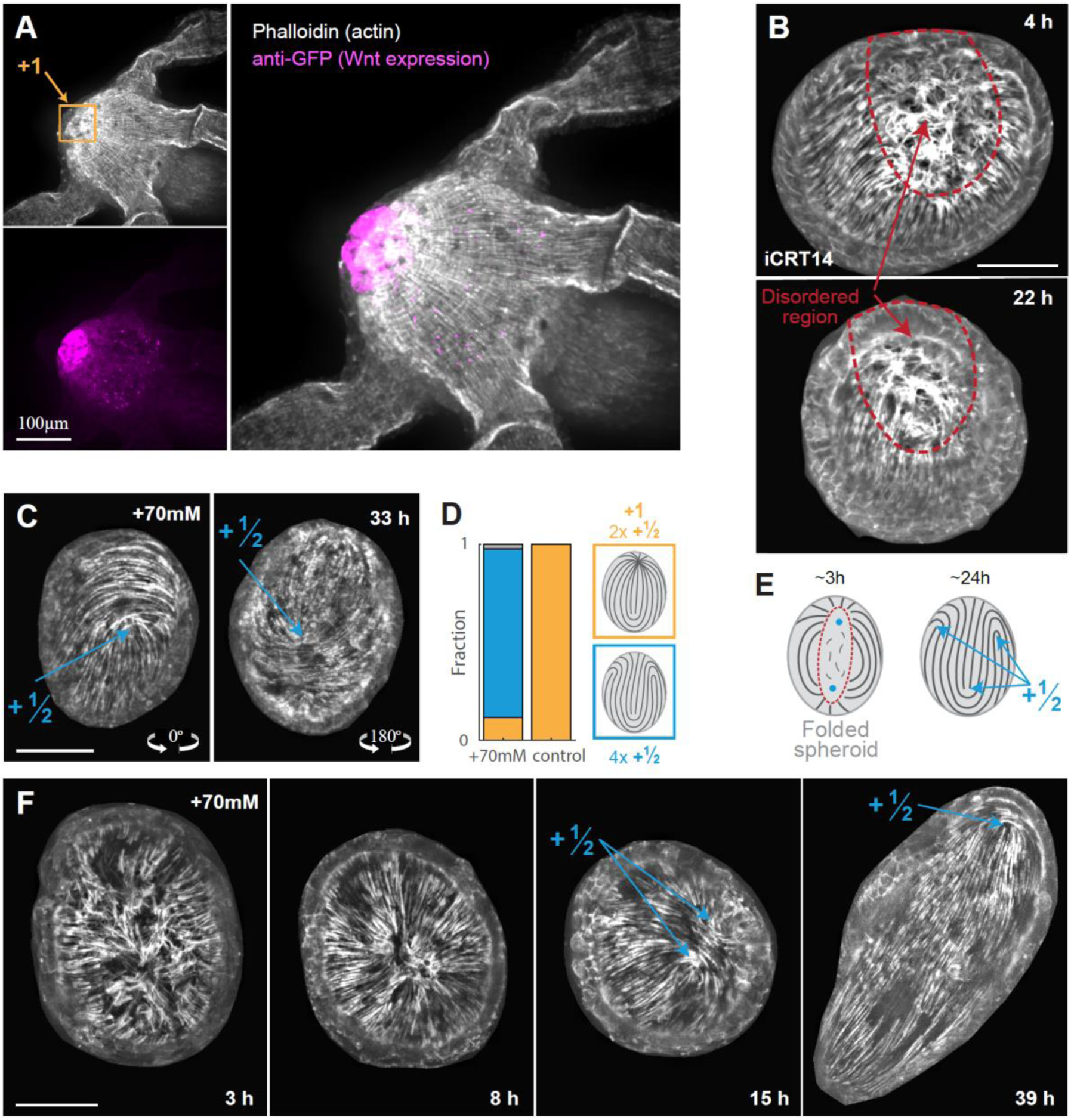
Perturbations of Wnt signaling and their influence on actin fiber dynamics. (A) Images of the head region in a mature Hydra. Transgenic Hydra from a Wnt-expression reporter strain are fixed and stained with anti-GFP to visualize Wnt expression and with phalloidin to visualize actin fibers (Methods). The Wnt expression region coincides with the +1 defect site. (B) The effect of iCRT14 treatment on actin fiber organization (Methods). Images from a time-lapse movie of a regenerating fragment treated with 5μM iCRT14, taken at an early (top) and later (bottom) time point (Movie 9). While the inherited aligned fibers are maintained, new fibers do not form in the disordered region (red line) and point defects do not emerge. (C-F) The effect of elevated medium osmolarity (+70mM sucrose) on actin fiber organization. (C) Images from opposite sides showing a tissue spheroid in isotonic medium displaying four +½ defects. (D) Bar plot comparing the defect configurations observed in isotonic medium to control conditions after 24-48 hours. (E) Schematic illustration of the actin fiber organization in a tissue spheroid placed in isotonic medium. The actin fibers reorganize into a nematic array in the initially disordered region and form a four +½ defect configuration. (F) Images from a time-lapse movie in isotonic media (Movie 10) showing a disordered region that orders into a +1 defect (8h), that is unstable and subsequently unbinds into a +½ defect pair (15h). The tissue stabilizes in a four +½ defect configuration and elongates along the fibers’ direction (39h).

To investigate the influence of down-regulation of Wnt on actin organization and tissue mechanics we use iCRT14, which inhibits β-Catenin-TCF interaction and has been shown to suppress Wnt signaling and disrupt regeneration of bisected Hydra (Cazet et al., 2021; Gufler et al., 2018). We follow the dynamics of excised tissue fragments subject to iCRT14 treatment, and examine its effect on actin fiber organization and tissue mechanics. Animals are pretreated with 5μM iCRT14 for 2 hours prior to excision (Cazet et al., 2021), and the excised tissues are subsequently maintained at the same iCRT14 concentration (Methods). Excised fragments fold and seal as untreated samples, with a similar pattern of inherited ordered fibers and a large disordered region (Fig. 6B, top). However, while the fibers in the ordered region remain stable, there is no induction of order in the disordered region (Fig. 6B, bottom, Movie 9). Thus, unlike control tissues, the treated samples remain essentially in their initial configuration with an inherited ordered region and a large disordered domain. Notably, under these conditions, aster-shaped defects fail to form and the samples do not regenerate.

Recent experiments showed that placing regenerating Hydra in elevated osmolarity, among various possible effects, also leads to suppression of Wnt signaling (Ferenc et al., 2021). These experiments corroborated previous findings, showing that when regenerating tissue fragments are placed in media that is isotonic with their lumen (∼70 mOsm), osmotic inflations cease (Kucken et al., 2008). They further showed that under these conditions, Wnt is initially up-regulated in response to the wound, but its expression is not sustained and subsequently declines, and the excised tissues fail to regenerate. Here we follow the tissue deformations and actin fiber dynamics in spheroids placed in Hydra medium supplemented with 70 mM sucrose. Initially, the folded spheroids have an ordered region and a disordered domain (Fig. 6E, left), which organizes into an ordered nematic fiber array (Movie 10), as in untreated samples. However, under isotonic conditions, the spheroids develop an unusual defect configuration with four +½ defects (Fig. 6C,D), rather than the characteristic defect configuration with a stable +1 defect at the future head site and a +½ defect pair at the future foot (Maroudas-Sacks et al., 2021). While we occasionally observe the formation of a +1 defect under isotonic conditions, these defects are unstable and dissolve into a +½ defect pair (Fig. 6E; Movie 10).

The treated tissues still undergo some stretching events with mechanical strain focusing at actin foci, yet these are diminished compared to control samples (Fig. S5; Movie 10). In particular, we observe fewer tissue ruptures under isotonic conditions and large stretching events are not observed once the four +½ defect configuration develops. Interestingly, even though the tissues fail to regenerate, they still elongate along the fiber direction, such that the two +½ defect pairs localize at either end of the cylindrical tissue, rotated 90° from each other (Fig. 6E, Movie 10). Overall, these results highlight the inherent coupling between actin fiber organization and Wnt signaling, and demonstrate that the formation of aster-shaped +1 defects can be disrupted when Wnt expression is perturbed.

### Mechanochemical model of Hydra regeneration

Our results show that the actomyosin fiber organization leads to mechanical strain focusing at defect sites. Notably, the primary site of strain focusing can be identified from the onset of regeneration and coincides with the location of the emergence of a head organizer at the tip of the regenerating head. Given the coincidence of +1 defect formation and the emergence of a new head organizer, an obvious question that arises is whether these events are related. We propose a self-organized mechanism in which mechanical strain focusing at actin foci and the localized biochemical signaling required for establishing a new head organizer reinforce each other (Fig. 7A). Specifically, we suggest a closed-loop feedback in which strain localized at the actin foci induces morphogen production. Concurrently, regions of high morphogen concentration align fibers along morphogen gradients and stabilize aster-shaped +1 defects, further enhancing the local strain focusing at these sites.

**Figure 7.**
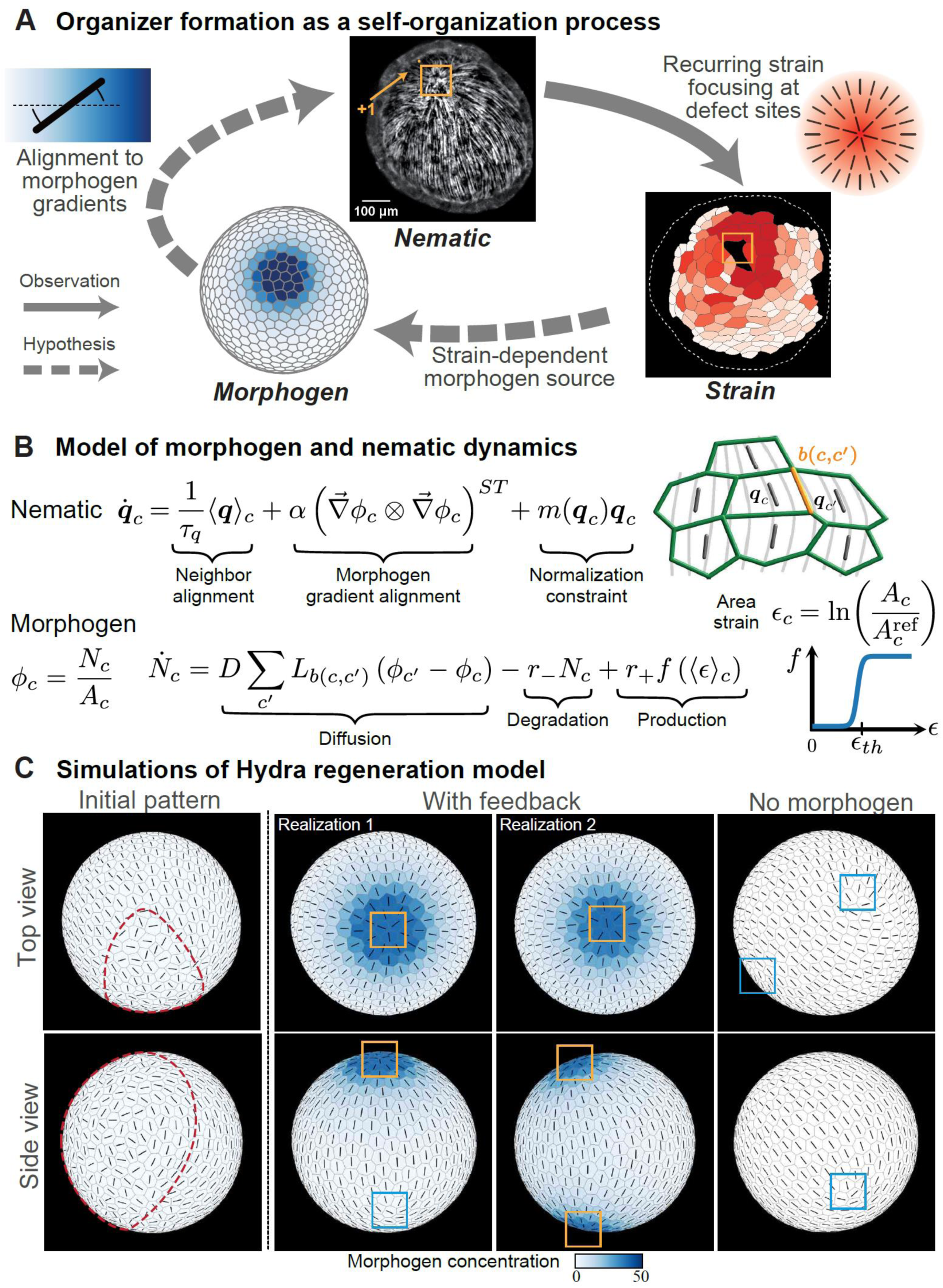
Mechanochemical model of Hydra regeneration. (A) Schematic illustration of the closed-loop feedback relating the nematic actomyosin fiber organization, tissue strain and a morphogen concentration field. Mechanical strain focusing induces local morphogen production. The fibers tend to align along the direction of morphogen gradients, enhancing the mechanical strain focusing at defect sites, and closing a positive feedback loop. (B) Equations describing the dynamics of the nematic and morphogen concentration fields in the model (Methods). The nematic field (**q_c_**) represents the local fiber orientation, which tends to align to fibers in neighboring cells and to local morphogen gradient. The morphogen diffuses between cells, is degraded at a constant rate and is produced in a strain-dependent manner described by *f*, where *ε*_*th*_ is the strain threshold. *N*_*c*_ is number of morphogen molecules in cell *c*, *φ*_*c*_ = *N*_*c*_/*A*_*c*_ is the morphogen concentration, *D* is an effective diffusion coefficient, *L*_*b*(*c*,*c*′)_ is length of the bond between cell *c* and its neighbor *c*^′^, and *r*_−_ and *r*_+_ are the morphogen degradation and production rates. The time-scale *τ*_*q*_ characterizes the nematic neighbor alignment rate, *m*(***q***_*c*_) is a Lagrange multiplier used to constrain the nematic magnitude to unity, and *α* characterizes the alignment of nematic to morphogen gradients. The recurring global contraction cycles are described by a time dependent active stress as in Fig. 5. (C) Simulation results showing regenerating tissues initiated with a partially ordered fiber configuration (as in 5B) and no morphogen (left panels; see SI for details). The tissues self-organize into a configuration with a +1 defect colocalized with a morphogen peak and a +½ defect pair, or into a configuration with two +1 defects colocalized with morphogen peaks (middle panels; Movies 11,12). Simulations with no morphogen production develop four +½ defects (right panels; Movie 13).

While the mechanistic details involved are unknown, recent results by us and others support different parts of the proposed feedback loop. The relation between mechanical strain and Wnt production was suggested by Ferenec *et al*., who correlated reduced tissue stretching under isotonic conditions with the inability to stabilize Wnt expression and regenerate a head (Ferenc et al., 2021). Note that in that work, the mechanical strain was assumed to be spatially homogenous, whereas our high-resolution imaging reveals strain focusing at the future organizer site. The idea that morphogen gradients can direct the orientation of actin fibers was recently suggested theoretically (Wang et al., 2023c), and is supported by observations showing that local Wnt expression can induce the formation of +1 defects in grafting experiments or during budding (Aufschnaiter et al., 2017; Wang et al., 2020). This notion is further supported by our current results showing the inability to stabilize aster-shaped +1 defects following perturbations of the Wnt pathway (Fig. 6).

We examine the feasibility of the proposed self-organization mechanism by developing a biophysical model of regenerating Hydra (Fig. 7; SI). This model extends the description of Hydra tissue mechanics (Fig. 5), by including the coupled dynamics of the nematic field and a biochemical morphogen throughout the regeneration process. The morphogen field is defined by specifying the number of molecules *N_c_*in each cell (Fig. 7B). We introduce a strain dependent morphogen source term as a sigmoid function of cell area strain, characterized by a threshold strain value ε_th_ and a saturation level *r_+_*(Fig. 7B, SI). The morphogen molecules can diffuse between cells and are degraded at a constant rate *r_−_*. The dynamics of the nematic field are governed by an interaction term that favors alignment of fibers in neighboring cells, as well as a coupling of fiber orientation to local morphogen gradients (Fig. 7B; SI). The recurring transient fiber contractions are emulated by periodically turning on the active stress in all cells with a period *T*_*ζ*_ and amplitude *ζ*_*M*_ using a Gaussian profile in time of width Δ*T*_*ζ*_.

Using this model, we simulate regeneration from tissue spheroids with an initial fiber configuration containing a large disordered region (as in Fig 5B) and no morphogen. We evolve the full dynamics of the model through a series of global contraction cycles, and examine whether the disordered region can reliably resolve to form a “head organizer”, i.e. a region containing a +1 nematic defect colocalized with a morphogen peak. We find model parameters that reproduce the emergence of a single organizer and a +½ defect pair on the opposite side (Figs. 7C, Movie 11; SI), as observed during Hydra regeneration. For the same set of model parameters, different initializations of the random nematic orientation in the disordered region can also yield a spheroid with two organizers (Fig. 7C, Movie 12). Thus, while the suggested feedback loop can robustly produce at least a single “head organizer”, it does not prevent the formation of two organizer sites. The outcome defect configurations depend on the strain threshold value, with two +1 defects being common at low values of ε_th_ and four +½ defects emerging at high threshold values (Fig. S6). As expected, that +1 defects are not stabilized in simulations with no morphogen production, and we obtain four +½ defects (Fig. 7C, Movie 13). Further work is needed to provide a detailed characterization of the regeneration outcomes as a function of model parameters.

## DISCUSSION

Our results show that the nematic fiber organization in folded spheroids obtained from excised tissue fragments contains two discrete actin foci. These foci develop into a characteristic set of topological defects, that eventually coincide with the sites of head/foot formation in the regenerated animal. As such, the location of the future head can be identified from the actomyosin fiber pattern at the onset of regeneration. We further show that these actin foci experience recurring mechanical strain focusing arising from transient contraction of the actomyosin fibers, converging at the defect core. This localized strain pattern provides a clear mechanical signature at the future head site, despite the presence of extensive disorder and fluctuations in the tissue.

Hydra has been a classic model system for studying axial patterning during animal morphogenesis for over a century, since the pioneering work of Ethel Brown who discovered the head organizer (Bode, 2012; Browne, 1909). Subsequent work established the presence of head-inducing activation and inhibition activities that are graded along the body axis (Shimizu, 2012). These observations led to the development of the famous Gierer-Meinhardt model that describes axial patterning in Hydra as a reaction-diffusion process (Meinhardt, 2012b). This model successfully integrated a large body of observations into a coherent picture that has dominated the field, in which tissue-scale gradients generate short-range activation and long-range inhibition of head formation. Despite its popularity, the mechanistic basis for this model is still unclear, and in particular, the nature of the source term, a crucial ingredient of the model, has remained obscure (Meinhardt, 2012a; Wang et al., 2023b). In the context of our work, it is also important to note that the Gierer-Meinhardt model does not consider the possible contribution of mechanics and the nematic fiber organization in the patterning process.

The characteristic defect pattern found in regenerating Hydra fragments, containing a +1 defect at the future head and a +½ defect pair at the future foot (Fig. 1), differs from the expected stable defect configuration for a nematic on a sphere with four +½ defects (Shin et al., 2008). Interestingly, we observe that spheroids placed in elevated osmolarity develop such a four +½ defect configuration (Fig. 6C,D). These results demonstrate that +1 defects are not a necessity of the nematic organization in Hydra, but rather require some additional stabilizing interaction. As previously suggested (Wang et al., 2023c), +1 defects can be stabilized through a coupling of the nematic field to morphogen gradients (Fig. 7). We hypothesize that this additional coupling is part of a mechanochemical feedback involved in head organizer formation.

Ferenc *et al*. recently suggested that tissue stretching due to osmotic inflations are essential for regeneration because of their role in stabilizing Wnt expression (Ferenc et al., 2021). While we believe that their results are valuable in highlighting the importance of mechanochemical coupling in Hydra regeneration, we disagree with some of their interpretations. In particular, while cycles of large osmotic inflations and ruptures typically occur during regeneration (Ferenc et al., 2021; Kucken et al., 2008), they are not essential (Fig. S7). Moreover, the localized Wnt expression at the future head was attributed to random symmetry breaking assuming homogenous tissue inflation. However, the observed tissue stretching is inhomogeneous, and the location of the future head is specified by the actin foci from the onset of regeneration.

We suggest a closed-loop positive feedback that couples topological defects in the nematic field with localized morphogen production, via the spatiotemporal strain patterns generated during actomyosin fiber contraction (Fig. 7). Within our model, the emergence of a head organizer at the defect site is not a coincidence or a corollary of biochemical patterning. Rather, we suggest that head organizer formation is a dynamic self-organization process that involves an interplay between morphogen dynamics and the nematic field. This hypothesis is appealing, since it naturally provides a mechanism for inducing local organizer activity at defect sites. The nematic field robustly generates discrete foci that produce localized strain, and the slow dynamics of the morphogen source field in this picture, arise from the slowly-varying dynamics of the actin fiber pattern (Maroudas-Sacks et al., 2021).

Obviously the formation of a new organizer is not solely determined by this mechanochemical coupling. The Wnt signaling pathway has an autocatalytic regulatory network and exhibits non-linear dynamics that can generate localized production on their own (Nakamura et al., 2011). Moreover, the memory of polarity in regenerating tissues implies the presence of additional relevant fields (Javois et al., 1988; Shani-Zerbib et al., 2022). While our current model of morphogen field dynamics is too simplistic, we believe that the proposed mechanochemical coupling between Wnt signaling and the nematic field, plays a role in facilitating the robust yet highly flexible regeneration capabilities in Hydra. Interestingly, similar mechanochemical coupling has been shown to be important in regenerating jellyfish fragments, where reorganization of the muscle fibers into aster-shaped defects was shown to define structural landmarks for the formation of the Wnt signaling center at the future manubrium (Sinigaglia et al., 2020).

The proposed mechanochemical feedback seems particularly important in regeneration from excised tissue pieces. Unlike organizer grafting experiments or bud formation, where defect formation initiates in a region containing a pre-established morphogen peak (Hobmayer et al., 2000), regenerating fragments are not expected to possess a well-defined initial biochemical pattern (Suzuki et al., 2023). The excised tissue spans only a small fraction of the body length of the parent animal and folds so that its originally head and foot-facing sides adhere to each other, making it unlikely that the folded spheroids contain a localized morphogen peak emanating from gradients in the parent animal. Similarly, the possibility that the head formation site is specified by a localized wound response (as could be the case in bisected animals (Cazet et al., 2021; Gufler et al., 2018)) appears unlikely, since the excised tissue fragment is wounded from all directions and folds into a spheroid in which the closure region spans nearly half of the regenerating spheroid (Fig. 1). Nevertheless, our observations show that head formation occurs in a well-defined location that can be identified from the actin fiber pattern at the onset of regeneration. Our ability to specify this location and identify a unique mechanical signature there, highlights the strong coupling between mechanics and biochemical signaling associated with organizer formation.

In the context of regenerating fragments, we suggest that mechanical fields can play an instructive role in specifying the future head site (Fig. 7). However, importantly, the relation between the presence of nematic defects and the establishment of a head organizer is not a simple causal relation. The proposed mechanism involves a feedback loop coupling the morphogen field and the nematic field, where neither is upstream or downstream from the latter (Fig. 7A). We suggest that depending on the context, different aspects of the initial conditions can be more or less instructional in guiding the highly flexible yet robust regeneration process. The memory of polarity, even in small excised tissues (Javois et al., 1988; Shani-Zerbib et al., 2022), in this view, does not arise from a well-defined pre-pattern, but rather emerges dynamically from the amplification of initial biases that are present in the excised tissue.

This work is only an initial step in deciphering the mechanochemical coupling underlying axial patterning in Hydra. While our proposed mechanochemical model is consistent with various experimental observations and has attractive features, it is nonetheless still speculative. Future work is required to establish this model from a phenomenological point of view, by proving that it can predict the behavior of regenerating tissues from various initial conditions (Livshits et al., 2017) and under different perturbations (Maroudas-Sacks et al., 2024; Wang et al., 2020). From a mechanistic point of view, activation of contraction in the nematic actomyosin fibers can nicely account for the observed relation between the nematic field and the tissue strain field (Fig. 5), yet the relation between the strain field and morphogen production is still unclear.

Tissue stretching has been shown in other systems to activate biochemical signaling (Martino et al., 2018). In particular, the Hippo-YAP pathway is known to respond to a diverse set of mechanical cues (Panciera et al., 2017), and is coupled to the Wnt signaling pathway (Azzolin et al., 2014). Similarly, while the Wnt pathway has been shown to influence actin dynamics (Stamatakou et al., 2015), how this could lead to nucleation and alignment of myonemes along morphogen gradients remains unknown.

Our suggested mechanochemical model for Hydra morphogenesis bears similarities with emerging views of plant morphogenesis, where mechanical forces have been shown to play an instructional part in patterning through mechanochemical feedback that couple cytoskeletal alignment, mechanical stress/strain fields and the distribution of morphogens such as Auxin (Trinh et al., 2021). We believe that similar self-organized dynamics involving mechanochemical feedback between stresses generated by the actomyosin cytoskeleton and signaling pathways are broadly relevant for animal morphogenesis. More generally, the presence of feedback from the emerging structure back into the patterning process itself naturally provides developing systems with immense flexibility, while simultaneously promoting the emergence of robust outcomes with well-defined patterns.

## MATERIALS AND METHODS

### Hydra strains culturing and regeneration from tissue fragments

All the experiments are performed using transgenic Hydra strains expressing Lifeact-GFP in the ectoderm, generously provided by Prof. Bert Hobmayer (University of Innsbruck, Austria). Additionally, we use a Wnt expression reporter strain, HyWnt3:GFP-HyAct:dsRED transgenic strain (Nakamura et al., 2011; Vogg et al., 2019) }, kindly provided to us by Prof. Brigitte Galliot’s lab (Geneva University, Switzerland), for studying Wnt expression in fixed Hydra (Fig. 6A). Animals are cultured in standard Hydra culture medium (HM: 1mM NaHCO_3_, 1mM CaCl_2_, 0.1mM MgCl_2_, 0.1mM KCl, 1mM Tris-HCl pH 7.7) at 18° C. The animals are fed 3 times a week with live Artemia Nauplii and rinsed after 4-8 hours. Tissue fragments are excised from the middle body section of mature Hydra, ∼24 hours after feeding, using a scalpel equipped with a #15 blade. Tissue rings are excised by performing two nearby transverse cuts, and are subsequently cut into two to four parts by additional longitudinal cuts to obtain rectangular tissue pieces.

### Tissue labelling

To label specific tissue regions we use laser-induced activation of a caged dye (Abberior CAGE 552 NHS ester) that is electroporated uniformly into mature Hydra and subsequently uncaged in the desired region (Maroudas-Sacks et al., 2021; Shani-Zerbib et al., 2022). Electroporation of the probe into live Hydra is performed using a homemade electroporation setup. The electroporation chamber consists of a small Teflon well, with 2 perpendicular Platinum electrodes, spaced 2.5 mm apart, on both sides of the well. A single Hydra is placed in the chamber in 10μl of HM supplemented with 6-12mM of the caged dye. A 75 Volts electric pulse is applied for 35ms. The animal is then washed in HM and allowed to recover for several hours to 1 day prior to tissue excision. Following excision, the specific region of interest is activated by a UV laser in a LSM 710 laser scanning confocal microscope (Zeiss), using a 20× air objective (NA=0.8). The samples are first briefly imaged with a 30 mW 488 nm multiline argon laser at up to 1% power to visualize the Lifeact-GFP signal and identify the region of interest for activation. Photoactivation of the caged dye is done using a 10 mW 405nm laser at 100 %. The activation of the Abberior CAGE 552 is verified by imaging with a 10 mW 543 nm laser at 1%. Subsequent imaging of the Lifeact-GFP signal and the uncaged cytosolic label is done by spinning-disk confocal microscopy as described below.

### Fixation and immunofluorescence staining

Intact Hydra from the HyWnt3:GFP-HyAct:dsRED transgenic strain are relaxed in 2% urethane in HM for one minute, and fixed in 4% formaldehyde in HM for 1 hour at room temperature. Samples are permeabilized with 0.1% Triton-X100 in PBS (PBT; 3 washes X 5 min) and then incubated for 2 hours with 0.1 % Triton-X100, 3% Bovine serum albumin (BSA, w/v) in PBS (PBSBT). For staining, samples are incubated with an AlexaFluor 647-conjugated anti-GFP, rabbit polyclonal antibody (2 mg/ml; Invitrogen) and AlexaFluor 488-conjugated phalloidin (200u/ml; Invitrogen) diluted 1:1:100 in PBSBT, and incubated overnight at room temperature with gentle shaking. Subsequently, samples are washed three times in PBT (3 washes X 15 min) and then in PBS (3 washes X 5min). Each sample is then placed in a cube of 2% low gelling agarose (sigma) and the cube is placed on a coverslip with its head facing the objective in order to image the mouth region.

### Sample preparation

Time-lapse live imaging is performed either in HM or in soft gel (0.5% low gelling point agarose (Sigma) prepared in HM to reduce tissue movement during imaging. The general characteristics of the regeneration process in soft gel are similar to regeneration in aqueous media. Samples are made in 50mm glass-bottom petri dishes (Fluorodish), polymer coverslip 24-well plates (Ibidi µ -plate 24 well, tissue culture treated), or in custom-made sample holders with a coverslip bottom. The samples are placed in few-mm sized wells cast of 2% agarose (Sigma) prepared with HM. For experiments in gels, the regenerating tissues are placed in liquefied 0.5% low gelling agarose gel that is cool enough not to damage the samples (∼ 35°C), ∼ 3-6 hours after excision (to allow the tissue pieces to first fold into spheroids), and the gel subsequently solidifies around the tissue. We add a few mm of HM above the gel to ensure it does not dry out during imaging.

To image samples from all directions by spinning-disk confocal microscopy, regenerating tissues are loaded within soft gel into FEP tubes (which have a similar refractive index to the surrounding solution) with a square cross-section and an internal diameter of 2 mm (Adtech). As above, the samples are inserted in liquefied 0.5% low gelling agarose gel and positioned within the FEP tubes. During imaging, the tubes are manually rotated to each of the 4 flat facets of the square tube and secured using a homemade Teflon holder to keep them stationary at each orientation. Images from 4 directions are acquired at the specified time points.

Samples for the up-and-under spinning-disk confocal microscope are prepared as follows. When using two air objectives, the samples are sandwiched between two glass coverslips at a set distance apart that are sealed from the environment to prevent leaks and fluid evaporation. The samples are placed within 2% agarose wells, which are filled with 0.5% low gelling agarose gel around the samples to maintain an aqueous environment between the coverslips. When using an air objective from below and a dipping lens from above, the samples are placed in a 50mm glass-bottom petri dish (Fluorodish) equipped with a homemade Teflon ring with tubing to allow perfusion of media to prevent evaporation. The samples are placed in wells made from 2% agarose that are filled with 0.5% low gelling agarose gel. When using the gel, the samples are layered with 3-4 ml of media on top of the gel. On the microscope stage we connect the small tubes from the Teflon ring to a peristaltic pump (Ismatec) and slowly flow media over the samples.

Samples for the light-sheet microscope are loaded in liquefied 0.5% low gelling agarose gel into a ∼ 1 cm long cylindrical FEP tube with an internal diameter of 2.15 mm (Zeiss Z1 sample preparation kit) and positioned along the central axis of the tube. The imaging is done through the FEP tubing, which has a similar refractive index to the surrounding solution.

### Pharmacological and osmotic perturbations

Pharmacological inhibition of the Wnt pathway with iCRT14 is done as follows (Cazet et al., 2021). Parent animals are preincubated in 5µM iCRT14 (sigma #SML0203) in HM for 2 hours prior to excision. Tissue fragments are cut and left to seal for ∼3 hours in the same solution (5µM of iCRT14 in HM). Tissue spheroids are then placed in wells prepared from 2% agarose containing 5µM iCRT14, and embedded in 0.5% low gelling agarose gel containing 5µM of iCRT14, and further layered with 5µM of iCRT14 in HM. The media with iCRT14 is refreshed every 24hr.

Perturbations with isotonic media (Ferenc et al., 2021) are done as follows. Tissue fragments are excised and allowed to seal for 3 hours in normal HM. Tissue spheroids are then placed in wells prepared from 2% agarose containing 70mM Sucrose (sigma, #84097), and embedded in 0.5% low gelling agarose gel containing 70mM Sucrose, and further layered with HM containing 70mM Sucrose with constant perfusion.

### Microscopy

Spinning-disk confocal z-stacks are acquired on a spinning-disk confocal microscope (Intelligent Imaging Innovations) running Slidebook software. The Lifeact-GFP is excited using a 50mW 488nm laser and the activated Abberior CAGE 552 is excited using a 50mW 561nm laser. Images are acquired with an EM-CCD (QuantEM; Photometrix). Time-lapse movies of regenerating Hydra are taken using either a 10× air objective (NA=0.5), a 10x water objective (NA=0.45), or a 20x air objective (NA=0.75).

The up-and-under setup is a custom, double spinning-disk confocal microscope (Intelligent Imaging Innovations) running Slidebook software, which enables imaging of the sample from two opposing angles - above and below. The Lifeact-GFP is excited using a 50mW 488nm laser and the activated Abberior CAGE 552 is excited using a 50mW 561nm laser. Images are acquired with two sCMOS cameras (Andor Zyla 4.1). Time lapse movies of regenerating Hydra are taken using a 10x air objective (NA=0.5) or a 20x air objective (NA=0.75) from below, and a 10x air objective (NA=0.3) or a 20x dipping lens (NA=0.5) from above.

Light-sheet microscopy is done on a Lightsheet Z.1 microscope (Zeiss). The light-sheet is generated by a pair of 10x air objectives (NA=0.2), imaged through 20× water objectives (NA=1), and acquired using a pair of CMOS cameras (PCO.edge). The Lifeact-GFP is excited using a 50mW 488nm laser and the activated Abberior CAGE 552 is excited using a 50mW 561nm laser. 4 views from different angles are acquired for the same sample by rotating the specimen. The imaging is performed using the “pivot scan” setting to minimize imaging artefacts that introduce streaking in the direction of illumination, yet some remnants of the streaking artefacts are still apparent in the images.

All 3D stacks in the spinning-disc microscopy systems and light sheet microscope are acquired at room temperature, typically taken at 3μm z-intervals. The time resolution of the movies ranges from 20 seconds to 30 minutes depending on the experiment. Imaging is done using appropriate emission filters at room temperature.

Time-lapse epifluorescence and bright field movies of regenerating Hydra are recorded on a Zeiss Axio-Observer microscope with a 5× air objective (NA = 0.25), at room temperature. Images are acquired on a charge-coupled device (CCD) camera (CoolSnap, Photometrix), and illuminated with a Xenon lamp. Time lapse imaging begins ∼ 4.5 hours after excision. and continues for 3 days at a time interval of 10 minutes.

### Image processing and analysis

The tools used for image processing and analysis are based on a combination of custom-written code with adaptation and application of existing algorithms, written in MATLAB, Python and ImageJ, as detailed below.

#### Creating masks of tissue region

In order to define the image region for analysis, masks were generated for every image based on the maximum intensity projections of the Lifeact-GFP signal, using automatic thresholding in ImageJ (‘Li’ method), and performing morphological operations (erosion, dilation, hole-filling) in MATLAB to obtain masks that accurately mark the tissue region in the image. All subsequent analysis was performed for the regions within the tissue masks.

#### Surface detection and layer separation

The regenerating Hydra tissue spheroids consist of a bilayered epithelium surrounding a hollow cavity. The 2D apical and basal surfaces of the ectoderm are computationally identified in the 3D fluorescence z-stacks of the Lifeact-GFP signal in the ectoderm. The supracellular actin fibers reside on the basal surface of the ectoderm while the apical junctions marking the cell outlines are visible on the apical surface. 2D projected images of the basal and apical surfaces of the ectoderm are automatically extracted from the 3D spinning-disk z-stack acquired with a 3μm z-interval using the “Minimum Cost Z surface Projection” plugin in ImageJ (https://imagej.net/Minimum_Cost_Z_surface_Projection). The cost images are generated by pre-processing the original z-stacks using custom-code written in MATLAB. First, the signal from the ectoderm layer was manipulated to make it more homogeneous within the layer without increasing its thickness, by applying the built-in MATLAB anisotropic diffusion filter.

Subsequently, we employ a gradient filter to highlight the apical and basal surfaces as the top and bottom boundaries of the ectoderm layer. The apical and basal surfaces are determined using the minCost algorithm (Parameters used: Rescale xy: 0.5, rescale z: 1, min distance: 15μm, max distance: 45μm, max delta z: 1, two surfaces). The surfaces given by the minCost algorithm are then smoothed by applying an isotropic 3D Gaussian filter of width 1-3 pixels (after rescaling to an isotropic grid matching the resolution in the z-direction) and selecting the iso-surface with value 0.5.

#### Image projection

2D projected images of the ectodermal actin fibers that reside on the basal surface of the ectodermal layer and the apical junctions, which are visible on the apical surface are generated by extracting the relevant fluorescence signal from the 3D image stacks based on the smoothed basal and apical surfaces determined above, respectively. For each x-y position, we employ a Gaussian weight function in the z-direction with a sigma of 3μm, which is centered at the z-value determined from the smoothed surface. The resulting 2D projected images are further subject to contrast limited adaptive histogram equalization (CLAHE) with MATLAB’s “adapthisteq” function with Rayleigh distribution and a tile size of 26μm. 2D projection of the photoactivated dye are generated by taking a maximum intensity projection of the 3D image stacks in the z-region between the smoothed apical and basal surface determined above.

#### Analysis of nematic field, defect sites and their outcome

The local orientation of the supracellular ectodermal actin fibers is described by a director field, which is oriented along the mean orientation determined in a small region surrounding every point in space. The nematic director field is determined from the signal intensity gradients in the 2D projected images of the ectodermal actin fibers, as described in (Maroudas-Sacks et al., 2021). Analysis of defect sites and their outcome in the regenerated animal is performed manually by identifying and following defects over time in 2D projections on the basal ectodermal surface 3D from fluorescent time-lapse movies of LifAact-GFP in regenerating Hydra. The defect sites included in the analysis are those that can be reliably tracked throughout the regeneration process from their formation until the final morphological outcome.

#### Segmentation of cell images

The contrast-enhanced apical surface images are segmented using EPySeg, a machine-learning based segmentation tool (Aigouy et al., 2020), applying a custom trained model using hand-segmented images of cell junctions as the ground truth for training. The resulting output is a binary segmentation mask for each image marking all cell outlines. The automatic segmentation is manually corrected using ‘Tissue Analyzer’ (Aigouy et al., 2020), also used for manual segmentation of the training set. Based on the segmentation masks, two additional output images are created for each original image using custom-written python code: an image marking all cell vertices, and an image marking all cell bonds, with each bond uniquely numbered (matching the output format of ‘Tissue Analyzer’).

#### Analysis of cell morphologies from segmented images

The segmentation masks, vertices and bond images hold the information on cell geometries and neighbor relations in the projected 2D image. This information is extracted from the images using custom written MATLAB code, and stored in a relational database format matching that of TissueMiner software (Etournay et al., 2016), with some modifications as detailed here. We apply a geometrical correction to the projected data to account for the curvature of the surface on which the cells reside, using the height map of the apical surface (see above). Since cell size is small relative to surface curvature, we assume each cell to be roughly planar in the plane tangent to the surface at the location of the cell. To determine the tangent plane for each cell, we use the mean normal to the surface over a window of width 20μm around the cell center. For each cell, we project the full cell outline from the segmented images onto the plane normal to the mean normal of the cell and subsequently smooth the resulting outline using a 5-point moving average before calculating bond lengths and cell areas to avoid artefacts due to discrete pixel resolution. 3D coordinates for cell centers, vertices and bonds (both original and smoothed), as well as neighbor relations, are stored in the database.

To query the database, quantify and visualize the data, we use a custom-written object-oriented MATLAB platform. The full documentation can be found on the lab Git repository. All geometrical measures for cells are calculated using the cell outlines on the tangent planes. Cell geometrical measures that are calculated include area (area inside the cell’s smoothed outline), perimeter (length of the smoothed outline), and cell shape anisotropy **Q**, defined as in (Merkel et al., 2017). Briefly, the cell shape anisotropy tensor **Q** is calculated by dividing each cell into triangles that connect each pair of adjacent vertices with the cell centre, and then calculating for each triangle how it differs from an equilateral triangle aligned along a set reference axis to obtain the anisotropy tensor for each triangle. For each cell, **Q** is then calculated by taking an area-weighted average over the triangles that comprise it. To calculate cell shape anisotropy along the fiber orientation, the cell shape tensor **Q** is then projected along the direction of the cell’s fiber orientation. To calculate the cell shape anisotropy relative to the direction to the defect, we draw a line between the cell and defect in the 2D projected image, project this orientation to the cell plane, and project **Q** along this direction. Fiber orientation and coherence per cell are assigned by taking the mean orientation and coherence of all pixels within the cell outline. The mean fiber orientation is projected to the cell’s tangent plane.

Cell’s distance from defects was determined based on the graph distance, which is the minimal number of cells connecting a given cell and the cell at the center of the defect.

To calculate the logarithmic area strain at a given distance from the defect, we subtract the mean logarithm of the cell areas of all cells at that distance before the event, from the mean logarithm of the cell areas of all cells at that distance during the peak of the event. This is mathematically equivalent to calculating the mean of the logarithm of the ratio of cell area during the peak and before the event for each cell. The last frame before the deformation was determined manually, and the peak of an event was determined as the frame in which the total area of the cells within a distance of 3 cells from the defect was largest.

#### Analysis of local deformation and rupture event statistics

Analysis of local tissue deformation and rupture events is performed manually by identifying frames within time-lapse movies in which these events occur, using the projected images of both the apical and basal surface of the ectoderm. Events are defined as coherent tissue deformation involving dozens of cells. We record all observed events and classify them according to the pattern of deformation and its localization. Since local stretching and ruptures typically occur simultaneously at both actin foci upon contraction, more than one event is typically recorded for a single contraction, describing the deformations observed in the different tissue regions. The categories used to classify events are as follows: small local stretching (cell area increase of less than 2-fold), large local stretching (cell area change of 2-fold or more), small ruptures, large ruptures (hole diameter ∼ tissue diameter), global contraction (large cell deformation in the direction perpendicular to fibers in the ordered region between the foci), and other cell shape deformations not involving area change (characterizes cell shape changes at 2x½ defect sites during global contractions). The criteria for defining a hole include a transient, visible gap in fluorescence in both the Lifeact-GFP signal and an additional tissue label if present, a change in tissue volume before and after the hole appearance and/or visible expulsion of tissue debris from the hole. Timing and location of the observed deformation and rupture events are recorded, and the sites are followed over time throughout the regeneration process (using fiduciary landmarks in the form of local photoactivated fluorescent tissue labels) to determine the fiber organization and morphological outcomes at these sites. To obtain statistics for the cumulative event distribution over time, time-lapse movies are taken in the up-and-under system to simultaneously image the samples from above and below with a time resolution of 2-2.5 minutes, which is shorter than the characteristic duration of an event (Fig. 2F,I).

#### Model Simulations

In our custom vertex model of Hydra tissue mechanics, individual cells are defined by the position of their vertices. Since these vertices are in general not all in the same plane, we need to define the area and local tangent plane for each cell (Fig. S8A). To this end, we first determine the cell center, 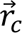, as the mean position of all the cell vertices. We then define a collection of cell triangles that each contain two neighboring vertices and the cell center. The cell area is defined as the sum of all cell triangle areas. The normal vector to the cell tangent plane is defined as the area weighted average of all the cell triangle normal vectors (see SI for more details).

The vertex model energy function is defined as,

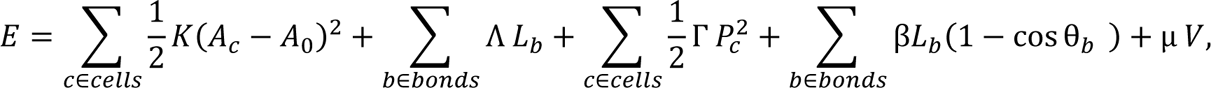

where *A*_*c*_ and *A*_0_ are the cell areas and preferred cell area, *K* is the area elastic modulus, Λ and *L*_*b*_ are the bond tension and bond length, Γ and *P*_*c*_ are the perimeter elastic modulus and cell perimeter, *β* is the cell-cell bending modulus and *θ*_*b*_ is the angle between normal vectors of neighboring cells that share the bond *b*. The total volume of the lumen is denoted by *V*, and *μ* is a Lagrange multiplier that ensures lumen incompressibility.

A vertex position, 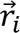, in the cellular network follows overdamped dynamics driven by the forces stemming from the gradient of the energy function 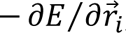, and an active force 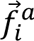 generated by the active stresses in all the cells to which this vertex belongs,

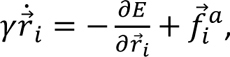

where *γ* is a friction coefficient. The active stress generated by a cell, 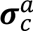, stems from the contraction of the actin fibers in that cell. We introduce a unit nematic tensor ***q***_*c*_ in the tangent plane of each cell, representing the average orientation of the actin fibers in that cell. When cells move or deform, the nematic tensor is convected and co-rotated with them (SI). We emulate global fiber contraction by simultaneously generating active stresses in all cells:

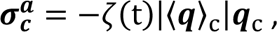

where the active stress magnitude *ζ*(*t*) is a Gaussian function of width Δ*T*_*ζ*_ that is centered at the time of the global fiber contraction and has an amplitude *ζ*_*M*_.

The cellular network evolves according to the forces acting on the vertices. If at any time-step of the simulation, the length of any bond becomes smaller than a threshold value 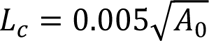 the two vertices of that bond are merged into a single vertex and the bond disappears. In this way 4-fold vertices are formed from merging of two 3-fold vertices. Such 4-fold vertices can resolve back into pairs of 3-fold vertices in one of two ways: reverting back to the original cellular configuration or rearranging the cells. A cell rearrangement event, also called a T1 transition, consists of the disappearance of one cell bond by the transient formation of a 4-fold vertex, that subsequently resolves into a new bond shared by cells that were previously not in contact. The stability of a 4-fold vertex is tested with respect to each of these two possible resolutions (SI). Note that our model supports vertices with an arbitrary large number of bonds, but in practice we observe only 4-fold vertices in our simulations.

To simulate the focusing of isotropic strain during stretching events (Fig. 5), we use a fixed nematic pattern that emulates the experimentally observed pattern of fiber alignment (Fig. 5B,C,left panels), and follow the vertex dynamics through global fiber contraction activation. Since we do not observe cell rearrangements in the Hydra ectoderm during stretching events, we chose parameters of the vertex model for which the cellular network has a negative two-dimensional Poisson ratio (see SI). This allows us to generate large isotropic strains without inducing cell rearrangements. Furthermore, we take the cell active stress amplitude to be equal to 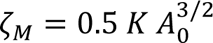, such that the isotropic strain profile we obtain in the simulation of a single event is comparable to the one experimentally observed in Hydra (Figs. 3, S4).

In order to study the proposed mechanochemical feedback between the nematic actomyosin fiber organization, mechanical strain in the tissue and morphogen concentration (Fig. 7A), we consider the coupled dynamics of the nematic field and a morphogen concentration field (Fig. 7B). The amount of morphogen in a cell *c* is described by the number of molecules *N*_*c*_ and develops over time according to,

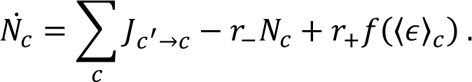

The first term reflects the contribution of diffusive fluxes between neighboring cells, that are of the form *Jc*′→*c* = *DL_b_*(*c*,*c*′)(*φc*′ − *φ_c_*), where *D* is an effective diffusion coefficient that characterizes inter-cellular diffusion, *L_b_*(*c*,*c*′), is the length of the bond shared by cell *c* and its neighbor *c*^′^, and *φ*_*c*_ = *N*_*c*_/*A*_*c*_ is the average morphogen concentration inside the cell. In this model intra-cellular diffusion is assumed to be much faster than transport between cells, so the morphogen concentration at the interface between two cells can be approximated by the average cellular concentration, *φ*_*c*_. Furthermore, we assume that the morphogen is degraded at a constant rate *r*_−_, and produced at a rate that is an increasing function of the cell area strain 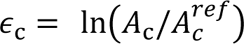, where 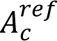 is the initial cell area at t=0. We use a sigmoid production function *f*(ɛ_*c*_) = 1/2 tanh(*w*(ɛ_c_ − *ε*_*th*_)/ɛ_*th*_), with *w* = 10, so that *f* increases sharply as a function of the strain around the threshold strain value ɛ_*th*_. Therefore, the production is small below ɛ_*th*_ and quickly saturates to *r*_+_ above it. Note that, without loss of generality, we can set *r*_+_ = 1, which corresponds to expressing *N*_*c*_ in units of *r*_+_/*r*_−_. The morphogen dynamics equation is therefore characterized by a degradation time-scale τ_−_ = 1/*r*_−_ and a length-scale 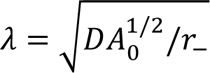, which reflects the length scale a morphogen diffuses before degrading.

To describe dynamics of the nematic field, we assume that the nematic orientation in each cell is coupled to the nematic orientation in neighboring cells, and we further introduce a coupling of the nematic to the local morphogen concentration gradient,

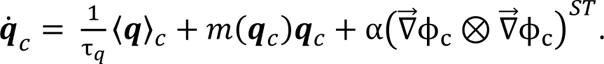

Here, the neighbor alignment is introduced through the average neighbor nematic operator ⟨***q***⟩_*c*_. On a flat surface, the average neighbor alignment would simply correspond to the arithmetic mean of the nematic tensors in neighboring cells. However, on a curved surface we need to account not only for in-plane alignment but also for the effect of surface curvature. In particular, since the nematic in each neighboring cell, ***q***_c′_, is constrained to the tangent plane of that cell, we first determine the projection ***P***_*c*_(***q*_*c*_**′) to the tangent plane of cell *c* (Fig. S8C), and then determine ⟨***q***⟩_*c*_ = ∑_*c*_′ ***P***_*c*_(*q*_*c*_′) /*n*_*c*_, where *n*_*c*_ is the number of neighboring cells. The dynamics of neighbor alignment is characterized by a time-scale τ_*q*_. We consider a nematic field with a fixed magnitude, describing the average orientation of actin fibers in each cell. The nematic magnitude is maintained equal to 1 through a Lagrange multiplier *m*(*q*_*c*_).

To describe the alignment of the nematic field to the local morphogen gradient we first have to calculate the gradient on the curved surface. As mentioned above, we assume that the intra-cellular diffusion is fast compared to inter-cellular diffusion. To estimate the overall morphogen gradient across a cell we have to take into account the morphogen concentrations in neighboring cells (Fig. S8D). For each cell we aim to find a gradient vector 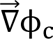 that satisfies relations 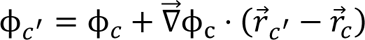 for all neighboring cell *c*^′^. However, in general these relations cannot all be satisfied with a single vector and, so we define the morphogen gradient to be the vector 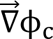 that produces the least-squared error of these relations (see SI for technical details). Finally, due to the nematic symmetry, the nematic will align only to the axis of the morphogen gradient, independent of the polarity of the gradient along this axis. For this reason, the nematic aligns to a tensor constructed as the traceless-symmetric component of the outer product 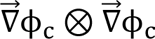. The parameter α characterizes the strength of alignment of the nematic to the morphogen gradient.

The choice for all the model parameter values used in the simulations is given in Table S1, and the rationale behind these choices is presented in the SI.

## ACKNOWLEDGEMENTS

We thank Erez Braun for many valuable discussions and comments on the manuscript. We thank Gidi Ben-Yoseph for excellent technical help and Lital Shani-Zerbib for her contributions to the development of the image analysis pipeline. We thank Alexandra Schauer, Jana Fuhrmann, Alex Mogilner, Aurelien Roux and Ram Adar for their comments on the manuscript. We thank Ben Atkinson from Intelligent Imaging Innovations for his help in designing the up-and-under spinning disk microscopy system, Nitzan Dahan from the LS&E Imaging Center for advice and help in imaging, Eran Kafri for help in 3D visualization, and the participants of the UCSB 2023 QBio course and the KITP program for discussions and providing a stimulating environment. This work was supported by a grant from the European Research Council (ERC-2018-COG grant 819174) to KK.

KK dedicates this work to the memory of Naomi Lindenstrauss.

