## Supplementary Information for "Mechanical strain focusing at topological defect sites in regenerating Hydra"

### I. MODEL OF REGENERATING HYDRA

Here we describe the vertex model of a regenerating Hydra that we developed to study the dynamics of Hydra tissue during the early stages of regeneration. Our model describes a tissue spheroid that encloses an incompressible fluid-filled lumen. The mechanics of the tissue is defined by an energy function that accounts for tissue elasticity. Furthermore, our model contains a nematic tensor field and a scalar morphogen field that are coarse-grained on the scale of individual cells and represented by a nematic tensor and a morphogen concentration value in each cell. We first employ this model to study how active stresses generated by the nematic field lead to focusing of area strain at locations of nematic +1 defects. Then, we incorporate a positive feedback-loop between tissue mechanics, nematic defects and a diffusing morphogen and show that it can provide a robust mechanism for +1 nematic defect stabilisation, co-localised with a morphogen source.

#### A. Vertex model of Hydra body mechanics

We model the mechanics of regenerating Hydra tissues using a custom vertex model of a curved deformable surface. We represent the Hydra myoepithelial bilayer by a single layer of cells that encloses a lumen. Cells are defined by a closed sequence of straight line segments, called cell bonds. Each bond is shared by two neighboring cells and its end-points are called vertices. Vertices, at which three or more bonds meet, are the degrees of freedom defining the state of the cellular network (Fig. S8A). The cellular network topology is defined by cell neighborship relations, which can change by cell rearrangements, also called T1 transitions [1].

In our model, cell bonds do not yet uniquely define a cell surface, as all the bonds of a given cell will in general not lie in the same plane (Fig. S8A). To define the surface of a cell $c$ we first define its center to be the mean position of all its vertices $i$

$$\vec{R}_c = \frac{1}{N_c}\sum_{i\in c}\vec{r}_i, \tag{1}$$

where $N_c$ is the number of cell vertices. We define the cell surface as a set of triangles generated by connecting each cell bond with the cell center by straight line segments. For each such triangle we define the triangle normal vector

$$\hat{n}_{c,i} = \frac{(\vec{r}_i - \vec{R}_c)\times(\vec{r}_{i+1} - \vec{R}_c)}{\left|(\vec{r}_i - \vec{R}_c)\times(\vec{r}_{i+1} - \vec{R}_c)\right|}, \tag{2}$$

where the vertices $i$ and $i+1$ are assigned such that the normal vector points outward, relative to the enclosed lumen. We use this convention in the rest of this Supplemental Material. The cell area is calculated as the sum of the triangle areas,

$$A_c = \frac{1}{2}\sum_{i\in c}|(\vec{r}_i - \vec{R}_c)\times(\vec{r}_{i+1} - \vec{R}_c)|. \tag{3}$$

We define a tangent plane for each cell by calculating a cell normal vector $\hat{n}_c$ which corresponds to the area weighted average of the normal vectors of all the cell triangles. In each cell $c$, we define a cell coordinate system $\mathbf{O}_c$ centered at the cell center $\vec{R}_c$, and the $z-$axis of the coordinate system coincides with the cell normal vertor $\hat{n}_c$.

The volume of the lumen enclosed by the cellular network is evaluated as the sum of tetrahedral volume elements

$$V_{\rm lumen} = \frac{1}{6}\sum_{c\in\text{cells}}\sum_{i\in c}\left[(\vec{r}_i - \vec{R}_c)\times(\vec{r}_{i+1} - \vec{R}_c)\right]\cdot\vec{r}_i \tag{4}$$

We define the mechanical energy of the cellular network by incorporating standard vertex model terms; cell area elasticity, bond tension, bond perimeter elasticity and an additional term that introduces a bending energy to reflect the energetic cost of bending a tissue of finite thickness,

$$E_0(\{\vec{r}_v\}) = \sum_{c\in\text{cells}} \frac{K_c}{2}(A_c - A_{c,0})^2 + \sum_{b\in\text{bonds}} \Lambda_b L_b + \sum_{c\in\text{cells}} \frac{\Gamma_c}{2} P_c^2 + \sum_{b\in\text{bonds}} \beta_b L_b (1 - \hat{n}_{b,c}\cdot\hat{n}_{b,c'}) \quad .$$

Here, $K_c$ and $A_{c,0}$ are the area elastic modulus and preferred cell area, $\Lambda_b$ and $L_b$ are bond tension and length of bond $b$, $\Gamma_c$ and $P_c$ are perimeter elastic constant and perimeter of cell $c$, $\beta_b$ is the bending elastic modulus and $\hat{n}_{b,c}$ , $\hat{n}_{b,c'}$ are, respectively, the normal vectors of cells $c, c'$ that share the bond $b$ and their product can be expressed as $\hat{n}_{b,c}\cdot\hat{n}_{b,c'} = \cos\theta_b$, where $\theta_b$ is the angle between the normal vectors. In the simulations of Hydra regeneration we use a constant value (i.e. a cell or bond independent value) for the following parameters: $K_c = K$, $A_{c,0} = A_0$, $\Lambda_c = \Lambda$, $\Gamma_c = \Gamma$, and $\beta_b = \beta$. For the numerical values of all the parameters used in the simulations see Table 1.

We further need to account for the fact that the fluid enclosed by the cellular network is incompressible and therefore its volume $V$ cannot be changed by the mechanical forces exerted by the cellular network. To implement this volume constraint we introduce a Lagrange multiplier $\mu$. Note that this method allows us to change $V$ in time to represent the changing lumen volume due to osmotic fluid influx and occasional flud eflux following tissue ruptures. However, for simplicity, in this work we keep $V$ constant over time. The mechanical energy function becomes

$$E(\{\vec{r}_v\}) = E_0(\{\vec{r}_v\}) + \mu V(\{\vec{r}_v\}). \tag{5}$$

The energy gradient generates a force on each vertex

$$\vec{F}_v = -\frac{\partial E(\{\vec{r}_v\})}{\partial \vec{r}_v}, \tag{6}$$

which we evaluate analytically for each of the terms in the energy function $E(\{\vec{r}_v\})$.

In order to describe the nematic pattern of actin fibers we introduce a nematic field on the surface of the cellular network represented in a coarse-grained manner by assigning a two-dimensional nematic tensor $\boldsymbol{q}_c$ to each cell. The nematic tensor $\boldsymbol{q}_c$ lies in the cell tangent plane.

The myoepithelial actin fibers exert mechanical stress upon activation [2]. In our model we introduce a nematic active stress in each cell which is proportional to the average cell nematic tensor in the cell and its neighbors $\boldsymbol{\sigma}_c^a = \zeta_c |\langle \boldsymbol{q}\rangle_c| \boldsymbol{q}_c$. The activity parameter $\zeta_c$ reflects the activation level of cell $c$ , and can vary from cell to cell depending on depending on the activation pattern in the tissue. Here, $|\langle \boldsymbol{q}\rangle_c|$ is the magnitude of the nematic tensor obtained as the average of the cell $c$ nematic and projected nematics of neighboring cells to the tangent plane of cell $c$ (Fig. S8C). In this way the norm of the active stress is reduced when nematics in neigboring cells are not fully aligned.

We translate the active stress generated by each cell $c$ to the corresponding forces on the vertices $\vec{f}^a_{c,v}$ of that cell using the active stress scheme from Ref. [3], generalised to three dimensions. The total active force on each vertex is the sum of the active forces from each of its cells $\vec{f}^a_v = \sum_{c(v)} \vec{f}^a_{c,v}$.

The dynamics of the cellular network follows the overdamped dynamical equation for the vertex positions,

$$\gamma \frac{d\vec{r}_v}{dt} = \vec{F}_v + \vec{f}^a_v \tag{7}$$

where $\gamma$ is a dissipation coefficient. In addition to movement of vertices, changes in the state of the cellular network can also occur through cell rearrangements during which cells exchange neighborship relations. In the vertex model such changes correspond to T1 transitions during which a bond between cells is lost and/or created. In our model, we implement T1 transitions so that whenever a bond becomes smaller than a threshold value $\epsilon_{T_1} = 0.005\sqrt{A_0}$, the bond is removed and its two vertices are merged into a single vertex. For example, merging of two three-fold vertices gives rise to a single four-fold vertex. If at any time a multi-fold (four or higher) vertex exists in the network we perform a check to test whether a new bond is energetically favourable, compared to retaining the multi-fold vertex. In particular, we consider all possible topological splittings of the multi-fold vertex into two vertices $i$ and $j$, while keeping the length of the new bond at zero length. For each of these topologies $(i, j)$, we calculate the force acting to extend the bond length: $f_{ij} = (1/2)|\vec{F}_i + \vec{f}^a_i - \vec{F}_j - \vec{f}^a_j| - (\Lambda_b + \Gamma P_{c(i)} + \Gamma P_{c(j)})$ where $i$ and $j$ are the two vertices of the new bond, $\vec{F}_i$ and $\vec{f}^a_i$ are the forces acting on vertex $i$ as described above, and $c(i)$ and $c(j)$ are the two cells that share the newly opened bond. We find the maximal value of $f_{ij}$ among all possible splittings $(i, j)$ and, if it is positive, we generate that bond. Then, we set the length of the new bond to $l_0 = 10\epsilon_{T1}$ along the axis of the force difference $\vec{F}_i + \vec{f}^a_i - \vec{F}_j - \vec{f}^a_j$. If the maximal $f$ is negative, we retain the multi-fold vertex.

#### B. Morphogen field

We introduce a morphogen field to our model of Hydra. The morphogen field is thought to describe an activator associated with the formation of the head organiser region [4]. We define the number of morphogen molecules $N_c$ in each cell $c$. The dynamics of the morphogen field is specified by a production-diffusion-degradation equation as follows,

$$\frac{dN_c}{dt} = D \sum_{c'} L_{b(c,c')} h_{b(c,c')} \left( \frac{N_{c'}}{V_{c'}} - \frac{N_c}{V_c} \right) - r_- N_c + r_+ f(\langle \epsilon \rangle_c), \tag{8}$$

where the first term corresponds to diffusive fluxes of morphogen $J_{c' \to c}$ from neighboring cells $c'$ to the cell $c$ (Fig. S8B), $L_{b(c,c')}$ and $h_{b(c,c')}$ are bond length and lateral interface height shared by cells $c$ and $c'$, and $D$ is the transport coefficient for morphogen diffusion between neighboring cells. $r_-$ is the degradation rate and $r_+$ is the production rate amplitude and $f(\langle \epsilon \rangle_c)$ is a sigmoid function that describes the morphogen production as a function of the cell area strain $\epsilon_c = \ln \left( A_c / A_c^{\mathrm{ref}} \right)$, where $A_c^{\mathrm{ref}} = A_c(t = 0)$ is the reference area, chosen as the area of cell $c$ at the start of the simulation, and $\langle ... \rangle_c$ indicates average over the cell $c$ and its neighbors. Specifically, we use $f(\epsilon_c) = \frac{1}{2} \left( \tanh(w(\langle \epsilon_c \rangle_c - \epsilon_{th})/\epsilon_{th}) + 1 \right)$ where $\epsilon_{th}$ is the strain threshold to produce morphogen and the parameter $w$ sets the range of strains over which the production is activated with increasing strain. In this work we use $w = 10$. Furthermore, we consider that diffusion inside cells is much faster than across cell boundaries so that the concentration inside a cell is approximately constant, compared to gradients across cells boundaries. Since the cell volume is not easily changed by mechanical deformation of cells we simplify Eq. 8 by introducing the 2D concentration $\phi_c = N_c / A_c$ so that our morphogen dynamics equation becomes

$$\frac{dN_c}{dt} = D \sum_{c'} L_{b(c,c')} \left( \phi_{c'} - \phi_c \right) - r_- N_c + r_+ f(\langle \epsilon \rangle_c) \tag{9}$$

where we have used the approximation that the height of cell-cell interfaces does not vary significantly across a single cell so that we can define a cell height $h_c$ which satisfies $A_c h_c = V_c$.

### C. Dynamics of the nematic field

The evolution of the nematic in each cell is given by

$$\frac{d\boldsymbol{q}_c}{dt} = \frac{1}{\tau_q} \langle \boldsymbol{q} \rangle_c + \alpha (\vec{\nabla} \phi_c \otimes \vec{\nabla} \phi_c)^{ST} + m(\boldsymbol{q}_c) \boldsymbol{q}_c, \tag{10}$$

where the first term is the standard alignment to the neighbor nematic orientation with a rate $1/\tau_q$. In the continuum limit on flat surfaces this term would correspond to the lowest term in the long-wavelength expansion of the nematic interaction energy in the one-constant approximation [5]. Here, we adjust the coupling to the curved surface by defining a neighbor-averaged nematic alignment as $\langle \boldsymbol{q} \rangle_c = \sum_{c' \in c} \boldsymbol{P}_c(\boldsymbol{q}_{c'}) / N_c$ where the cells $c'$ are neighbouring cell $c$, $\boldsymbol{P}_c(\boldsymbol{q}_{c'})$ is the projection of the nematic in cell $c'$ to the tangent plane of cell $c$ (Fig. S8C), and $N_c$ is the number of neighbours of cell $c$. The second term in Eq. 10 corresponds to nematic-morphogen interaction that favors alignment of the nematic to gradients of the morphogen field, with a coupling coefficient $\alpha$. Here the superscript $ST$ denotes the symmetric traceless part of the tensor, and $\vec{\nabla} \phi_c$ denotes the discrete gradient of the morphogen field at $\vec{r}_c$, determined as described below (Fig. S8D). The last term in Eq. 10 imposes the magnitude constraint $|\boldsymbol{q}_c|^2 = 1$.

We calculate the discrete gradient $\vec{\nabla} \phi_c$ of the morphogen field following the standard procedure based on least square optimization as follows. We define a cost function,

$$e_c = \sum_{c'} \left[ \phi_{c'} - \phi_c - \vec{\nabla} \phi_c \cdot \vec{r}_{c,c'} \right]^2 \tag{11}$$

where the sum is over cells $c'$ that are neighbors of cell $c$. To find the distance between neighboring cell centers we deterime the geodesic line connecming them along the surface (determined by the cell triangles), see Fig. S8D. Concretely, we define the displacement vector $\vec{r}_{c,c'}$ between neighboring cell centers by rotating each neighboring cell $c'$ of the cell $c$ around their shared bond $b(c, c')$ such that its center lies in the tangent plane of the cell $c$. The length of $\vec{r}_{c,c'}$ is equal to the distance between cell centers along the geodesic.

The discrete gradient $\vec{\nabla} \phi_c$ is then defined as the one that minimizes this cost function and is given by

$$\vec{\nabla} \phi_c = \boldsymbol{M}_c^{-1} \cdot \sum_{c'} (\phi_{c'} - \phi_c) \vec{r}_{c,c'}, \tag{12}$$

where

$$\boldsymbol{M}_c = \sum_{c'} \vec{r}_{c,c'} \otimes \vec{r}_{c,c'}. \tag{13}$$

#### 1. *Transport of the nematic*

As the cells move in space, the nematic $\boldsymbol{q}_c$ in each cell is convected and co-rotated with the cell. Nematic convection is implemented by maintaining the nematic position at the cell center as it moves. To implement the co-rotation of the nematic we devised a two step algorithm that accounts for out-of-plane rotations, as defined by changes of the cell normal vector, and then the remaining in-plane rotations of each cell: In the first step we determine the cell normal vector $\hat{n}_c$ at times $t$ and $t + \Delta t$, see Eq. 2. To make sure that the nematic remains in the cell tangent plane at the time $t + \Delta t$, we rotate the nematic by a rotation matrix $\boldsymbol{R}_1$ that corresponds to rotation between the two normal vectors: $\hat{n}_c(t + \Delta t) = \boldsymbol{R}_1 \hat{n}_c(t)$. In the second step we account for rotation of the cell within its tangent plane defined by the normal vector $\hat{n}_c(t + \Delta t)$. Following [6], we compute a rotation vector $\vec{\Omega}_c = \boldsymbol{I}_c^{-1} . \boldsymbol{L}_c$ where $\boldsymbol{I}_c = \sum_{v \in c} \left[ (\vec{r}_v - \vec{R}_c)^2 \boldsymbol{I}_3 - (\vec{r}_v - \vec{R}_c) \otimes (\vec{r}_v - \vec{R}_c) \right] / N_c$

is the moment of inertia of the cell and $\boldsymbol{L}_c = \sum_{v\in c}(\vec{r}_v - \vec{R}_c) \times (\vec{u}_v - \vec{U}_c)/N_c$ is the angular momentum of the cell. Here, $\vec{u}_v$ denotes the velocity of the vertex $v$, defined as $\vec{u}_v = (\vec{r}_v(t+\Delta t) - \vec{r}_v(t))/\Delta t$ and $\vec{U}_c$ denotes the velocity of the cell $c$ defined as $\vec{U}_c = \sum_{v\in c} \vec{u}_v/N_c$. Finally, the rotation vector is obtained by projecting $\vec{\Omega}_c$ to $\hat{n}_c(t+\Delta t)$, which determines the second rotation matrix $\boldsymbol{R}_2 = \boldsymbol{\mathcal{R}}(\vec{\Omega}_c[\vec{\Omega}_c \cdot \hat{n}_c(t+\Delta t)]/|\vec{\Omega}_c|)$, where $\boldsymbol{\mathcal{R}}$ is the operator converting a rotation vector to the corresponding rotation matrix. To complete the second step we rotate the nematic by $\boldsymbol{R}_2$. The local reference frame of each cell is also corotated in the same manner.

### II. MODEL PARAMETERS

Our model of Hydra tissue contains a number of parameters that we have to choose appropriately. Below, we explain the rationale behind this and Table 1 summarizes all the parameter values used. The mechanical parameters $K$, $\Gamma$, and $\Lambda$ were first constrained such that the cellular network is in the solid phase [7, 8], since the behavior of the Hydra during early regeneration appears to be elastic. Furthermore, following our observation that cell rearrangements are rare even during the large celluar stretching events, as well as motivated by recent experimental observations in other epithelial tissues [9, 10], we decided to set the parameters such that the two-dimensional Poisson ratio of the cellular network is negative. To this end we follow the analysis of the vertex model in [11] and set $\overline{\Lambda} = \Lambda/(KA_0^{3/2}) = 0$ and $\overline{\Gamma} = \Gamma/(KA_0) = 0.1$. We use $\sqrt{A_0}$ as length unit, and $KA_0^2$ as a unit of energy. Furthermore, we set the time unit in simulations so that the elastic cell relaxation timescale is $\overline{\tau} = KA_0/\gamma = 1$. Although we do not have a precise measurement in Hydra, we expect that this timescale corresponds to a value somewhere in the range of $1s$ - $1min$ in experiments. For the choice of bending modulus value $\beta = 0.1KA_0^{3/2}$ we did not have a constraint from literature or experiments so we chose an intermediate value that was sufficient to prevent strong bending at scale of single cell, but not too large as to dominate the tissue mechanics.

We chose the cell active stress amplitude to be $\zeta_M = 0.5\ K\ A_0^{3/2}$, such that the isotropic strain profile we obtain in the simulation of a single contraction event is comparable to the one experimentally observed in Hydra (Fig. 3, SI Fig. 5).

The parameters controlling degradation and diffusion of morphogen define a length-scale over which the morphogen diffuses from the source $\lambda = \sqrt{D/(A_0 r_-)}$. We take a value of $r_- = 0.0007/\overline{\tau}$. Motivated by observations of the decay time-scale of the 'head forming potential' in regenerating Hydra reported in Ref. [12] we take the corresponding time-scale $\tau_\phi = 1/r_-$ to be 10hr in experiments. This also sets the elastic cell relaxation timescale, introduced above, to be $25s$. We next choose the morphogen diffusion coefficient to be $D = 0.0063\sqrt{A_0}/\overline{\tau}$, such that $\lambda \approx 3\sqrt{A_0}$. Furthermore, the total amount of morphogen in the system $N_0$ is proportional to $r_+/r_-$. We now choose to measure molecule number in unit where $r_+ = 1/\overline{\tau}$ and $r_-$ is chosen as above. Therefore, in our simulations we set $r_+ = 1$.

In order to characterize how the model parameters influence the competition between the nematic neighbor alignment and alignment to morphogen gradients, we construct a dimensionless number that quantifies this competition. Assuming that the region in which morphogen is produced is small compared to the region over which it diffuses from the source, we can estimate the morphogen concentration amplitude as the ratio between the total amount of morphogen and the area over which the morphogen diffuses $A_d$, which is proportional to $\lambda^2$. Therefore, we expect the amplitude of the morphogen concentration to scale as $N_0/A_d = r_+/\left(r_-\lambda^2\right)$, and at distance $x$ from the morphogen source, the morphogen alignment term will scale as $\alpha\left((N_0/A_d)/\lambda\right)^2 e^{-2x/\lambda} \sim \alpha\left(r_+/\left(r_-\lambda^3\ \right)\right)^2 e^{-2x/\lambda} \sim \alpha r_+^2 r_-/\left(D^3 A_0^{3/2}\right) e^{-2x/\lambda}$. Now, we define a dimensionless function $S(x)$ as a ratio of morphogen alignment strength to the neighbor alignment rate $1/\tau_q$ at distance $x$ from the source:

$$S(x) = \frac{\tau_q \alpha r_- r_+^2}{D^3 A_0^{3/2}} e^{-2x/\lambda}. \tag{14}$$

In the region where $S(x) \ll 1$, nematic neighbor alignment dominates, while in the region where $S(x) \gg 1$ nematic dynamics is dominated by the alignment to the morphogen gradient. The length-scale $\Lambda$ that separates these two regimes can be identified by setting $S(\Lambda) = 1$, so that

$$\Lambda = \frac{\lambda}{2} \ln\left(\frac{D^3 A_0^{3/2}}{\tau_q \alpha r_+^2 r_-}\right). \tag{15}$$

Now, we can constrain the choice of remaining parameters $\tau_q$ and $\alpha$ by setting $\Lambda \approx 3.7\sqrt{A_0}$, which is comparable to $\lambda$ so that alignment to the morphogen gradient is dominant within few cells from the source, while neighbor alignment is dominant further away from the source. Finally, we choose $\tau_q$ to correspond to 13min in experiments, which determines uniquely the value for $\alpha$. Note that our choice of $\tau_q$ is such that $1/\tau_q$ is longer than duration of contraction events, which we will take to be $\Delta T_\zeta = 8\overline{\tau}$, corresponding to about 7min in experiments (Fig. 2).

Finally, we choose to simulate a single contraction event by turning on the active stress magnitude uniformly in all cells with,

$$\zeta(t) = \zeta_M \exp\left(-\left(\frac{t-t_M}{2\sigma_M}\right)^2\right) \tag{16}$$

The duration of the pulse $\Delta T_\zeta$, defined as the full width at half maximum $\Delta T_\zeta = 2.355\sigma_M$ is chosen to correspond

| Model parameter | Expression | Value | Value choice rationale |
|---|---|---|---|
| Cell length-scale | $\sqrt{A_0}$ | 1 | unit length |
| Cell energy-scale | $KA_0^2$ | 1 | unit energy |
| Elastic relaxation time-scale | $\overline{\tau} = KA_0/\gamma$ | 1 | unit time |
| Normalized bond tension | $\overline{\Lambda} = \Lambda/(KA_0^{3/2})$ | 0 | negative vertex model Poisson ratio |
| Normalized perimeter elasticity | $\overline{\Gamma} = \Gamma/(KA_0)$ | 0.1 | negative vertex model Poisson ratio |
| Normalized bending modulus | $\overline{\beta} = \beta/(KA_0^{3/2})$ | 0.1 | |
| Normalized active stress magnitude | $\overline{\zeta}_M = \zeta_M/(KA_0^{3/2})$ | 0.5 | reproduces experimentally observed strain |
| Time interval between activity pulses | $T_\zeta/\overline{\tau}$ | 160 | corresponds to about 1 hr |
| Duration of activity pulse (FWHM) | $\Delta T_\zeta/\overline{\tau}$ | 18.84 | corresponds to about 8min |
| Morphogen degradation rate | $r_-\overline{\tau}$ | 0.0007 | corresponds to 1/(10hr) |
| Morphogen production rate | $r_+\overline{\tau}$ | 1 | corresponds to about 1/(25s) |
| Morphogen diffusion coefficient | $D\overline{\tau}/\sqrt{A_0}$ | 0.0063 | diffusion length scale of $\sim 3$ cells |
| Nematic alignment rate | $\overline{\tau}/\tau_q$ | 0.03 | corresponds to 13min |
| Nematic-morphogen coupling strength | $\alpha\overline{\tau}/A_0^3$ | 0.001 | |
| Strain threshold for morphogen production | $\epsilon_{\text{th}}$ | 0.25 | |

TABLE 1. Values of the model parameters used in the simulations. We summarize the rationale behind the choice in the last column. Correspondences to real times are based on experimental observations. For the parameters $\beta$, $\tau_q$ and $\alpha$ we did not have a strong constraint from literature or experiments and the value choice was based on the simulation outcome. The strain threshold value was systematically varied (Fig. S6). The value reported in the table corresponds to the value in Fig. 7C.

to about 8min in experiments. When simulating multiple contraction events spacing between subsequent pulses $i$ and $i+1$: $T_\zeta = t_{M,i+1} - t_{M,i}$ is taken to correspond to 1hr in experiments.

We sumarize the chosen values of the model parameters in Table 1

### III. SIMULATIONS OF A REGENERATING HYDRA

#### A. Preparation of initial state

In order to perform simulations of global contraction activation events in regenerating Hydra, as well as the dynamics of the nematic organisation and morphogen distribution we first have to prepare a state of the tissue that will serve as an initial condition from which we evolve the dynamical equations outlined above. We prepare the initial cellular network configuration for all simulations in this manuscipt by first generating a spherical tiling. To this end we start with an icosahedron and we apply alternating 4 "truncate" and 3 "dual" operations, as defined by the Conway polyhedron notation [13]. After the last "truncate" operation we have a cellular network made of 812 cells. The number of cells is chosen to be similar to the value in experiments. We then randomize the network by introducing stochastic fluctuations of the bond tension parameter $\Lambda_b$ independently in each bond using

$$\frac{d\Lambda_b}{dt} = -k_\Lambda(\Lambda_b - \Lambda_0) + \sqrt{2k_\Lambda}\xi(t) \tag{17}$$

where the first term corresponds to relaxation of $\Lambda_b$ to a target value $\Lambda_0$ at a rate $k_\Lambda$, and the second term corresponds to fluctuations in $\Lambda_b$ with $\xi(t)$ denoting gaussian white noise: $\langle\xi(t)\xi(t')\rangle = 2D_\xi\delta(t-t')$. We choose $k_\Lambda = 1, \Lambda_0 = -0.15, D_\xi = 0.35$. We proceed to randomize the cellular network for $T_{\text{rand.}} = 100\overline{\tau}$. During the randomization process cells that become very small in area are extruded from the network and by the end of randomisation we have 804 cells remaining. Finally, we revert the bond tension value in all bonds to the uniform value we use in the actual simulations (Table 1), and minimize the vertex model energy to the nearest local minimum.

#### B. Strain focusing at nematic defects

We simulate a single global contraction activation event during Hydra regeneration to characterize the strain pattern induced by global contraction of the muscle fibers whose orientation is described by the nematic field. Since events last only several minutes (see main text and Fig. 3D), which is faster than the time-scale on which the nematic fiber pattern evolves [14], we do not consider here changes in the nematic pattern during a single event. We thus keep the nematic in each cell fixed, except for changes associated with convection and co-rotation of the cells as the tissue deforms.

We consider two nematic configurations: In the first case, we emulate the initial fiber pattern observed just after the tissue fragment folds into a spheroid, where a large part of the tissue surface is covered by an ordered nematic array without defects, which surrounds a disordered region of total nematic charge +2 (Fig. 1 B). In the disordered region, we take nematic in each cell to be oriented randomly in the cell tangent plane. Different realisations of disorder correspond to different sampling of the nematic orientations in the disordered region. In the second case, we consider a fully ordered nematic field with one $+1$ defect and a pair of $+1/2$ defects on the opposite side, emulating the nematic fiber pattern observed in regenerating Hydra around $10-20$ hours after excision (Fig 1 C). In both cases, we activate the contraction in all cells simultaneously, following Eq. 16.

#### C. Mechanochemical model of Hydra regeneration

We simulate the dynamics of the proposed feedback mechanims that involves a strain focusing during contraciton events, mechanosensitive morphogen production and nematic alignment with morphogen gradients. The simulation are intitiated from the configuration of cells desribed above, with a large disordered region in the nematic field with total charge $+2$, emulating the early organization in the folded Hydra spheroid. For 10 different realisations of the nematic disordered region, we evolve the system through 50 contraction events. We record and report the state of the system after 30 contraction cycles (each with a duration of $\sim 1hr$), which is comparable to the duration of Hydra regeneration.


[1] Denis L Weaire and Stefan Hutzler, *The physics of foams* (Oxford University Press, 1999).

[2] John R. Szymanski and Rafael Yuste, “Mapping the whole-body muscle activity of hydra vulgaris,” Current Biology **29**, 1807–1817.e3 (2019).

[3] Jordi Comelles, Soumya Ss, Linjie Lu, Emilie Le Maout, S Anvitha, Guillaume Salbreux, Frank J¨ ulicher, Mandar M Inamdar, and Daniel Riveline, “Epithelial colonies in vitro elongate through collective effects,” eLife **10**, e57730 (2021).

[4] Bert Hobmayer, Fabian Rentzsch, Kerstin Kuhn, Christoph M. Happel, Christoph Cramer Von Laue, Petra Snyder, Ute Rothbächer, and Thomas W. Holstein, “Wnt signalling molecules act in axis formation in the diploblastic metazoan hydra,” Nature **407**, 186–189 (2000).

[5] Pierre-Gilles De Gennes and Jacques Prost, *The physics of liquid crystals*, 83 (Oxford university press, 1993).

### Supplementary Figures

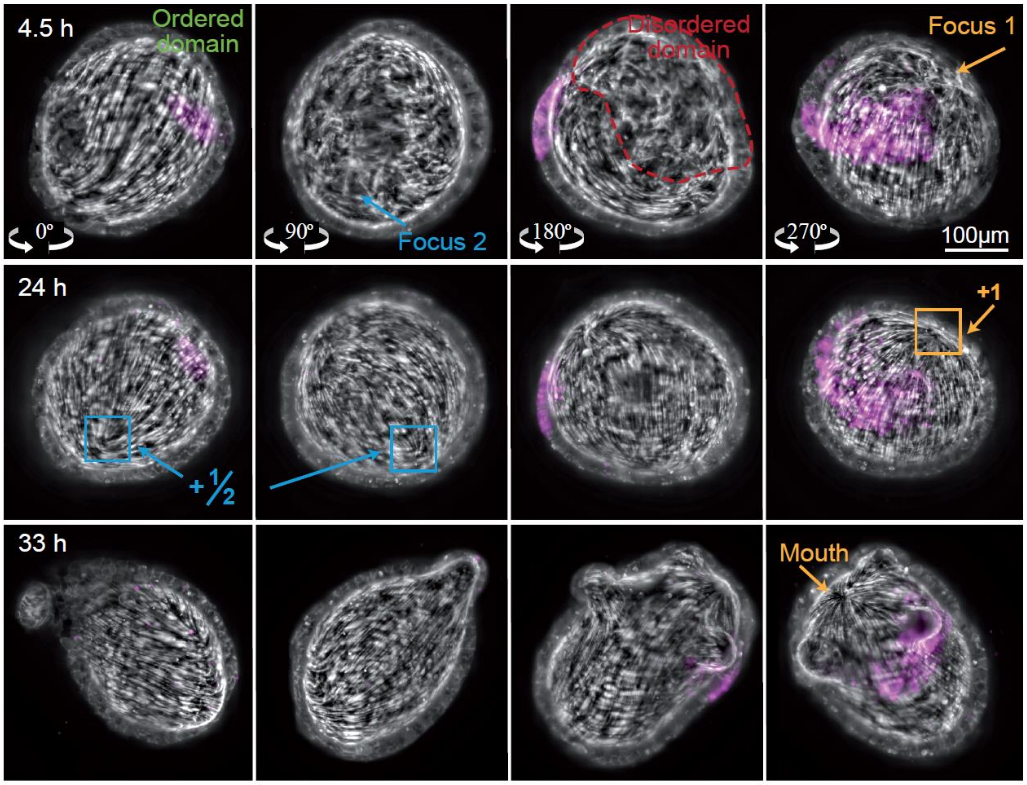


**Figure S1. 3D view of the actin fiber reorganization in a regenerating Hydra.** Images from a time-lapse light sheet movie of a regenerating Hydra expressing Lifeact-GFP in the ectoderm, showing projected views from four different directions at early (top), intermediate (middle) and late (bottom) stages of the regeneration process. The two actin fiber foci are visible in the folded spheroid (top panels), and develop, respectively, into an aster-shaped +1 defect at the future head site (orange labels), and a pair of +½ defects at the future foot site (cyan labels). The actin fiber organization (gray) is shown with an overlay the photoactivated dye (magenta; Abberior CAGE 552), identifying the same group of cells over time. All images are maximum projection images of the light sheet stack acquired from the direction indicated, showing the fiber organization at the basal surface of the ectoderm.

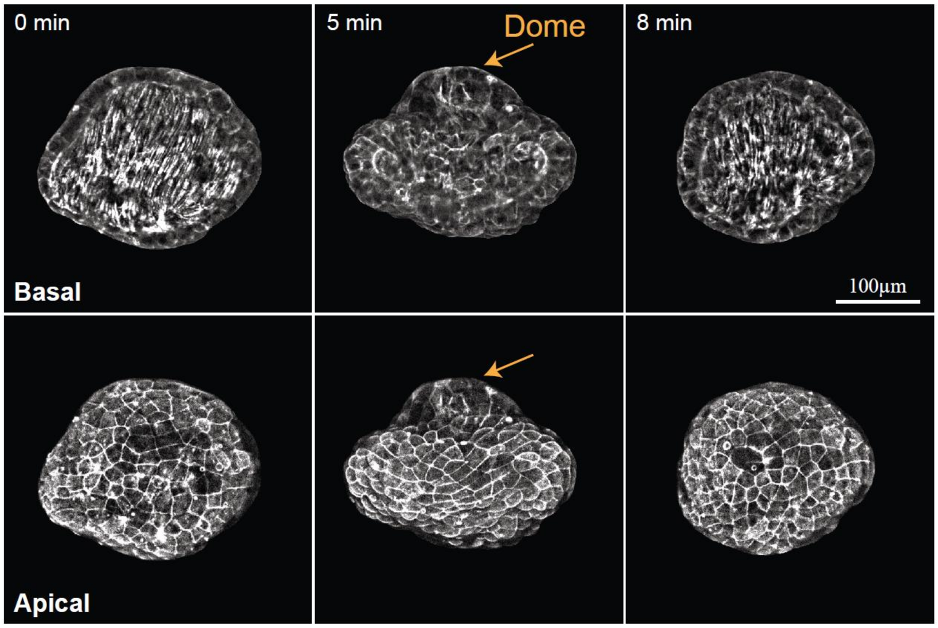


**Figure S2. Doming of the tissue during Hydra regeneration.** Images from a time-lapse spinning-disk movie of a regenerating Hydra expressing Lifeact-GFP in the ectoderm, exhibiting doming of the tissue at a defect site during a large tissue stretching event. The images show the fiber organization at the basal surface (upper panels) and the cellular organization at the apical surface (lower panels) before (left), during (middle) and after (right) the event. During the event, the tissue in the defect region undergoes localized stretching and exhibits an abrupt change in surface curvature resulting in the formation of a tissue dome (middle panels). Doming is suggestive of inhomogeneous or non-linear material properties that produce excessive stretching near the core of the event and induce out-of-plane deformations (Latorre et al., 2018).

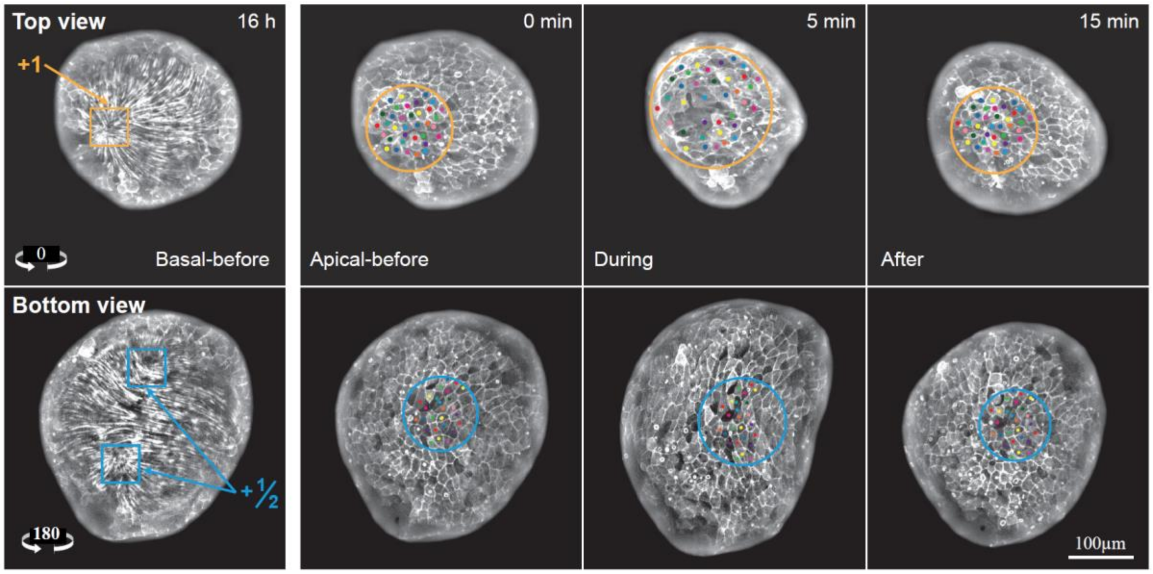


**Figure S3. Simultaneous mechanical strain focusing at both actin foci.** Images from a time-lapse up-and-under spinning-disk movie of a regenerating Hydra, during a global deformation event exhibiting mechanical strain focusing at both actin foci. The top view shows the deformations at and around a +1 defect, whereas the bottom view displays the deformations at a pair of +½ defects. For each view, the left image depicts the actin fibers at the basal surface of the ectoderm, and the right images show the cellular organization at the apical surface before (left), during (middle) and after (right) the event, with colored dots marking manually-tracked individual cells. All images are projected spinning-disk confocal images of regenerating tissues expressing Lifeact-GFP in the ectoderm.

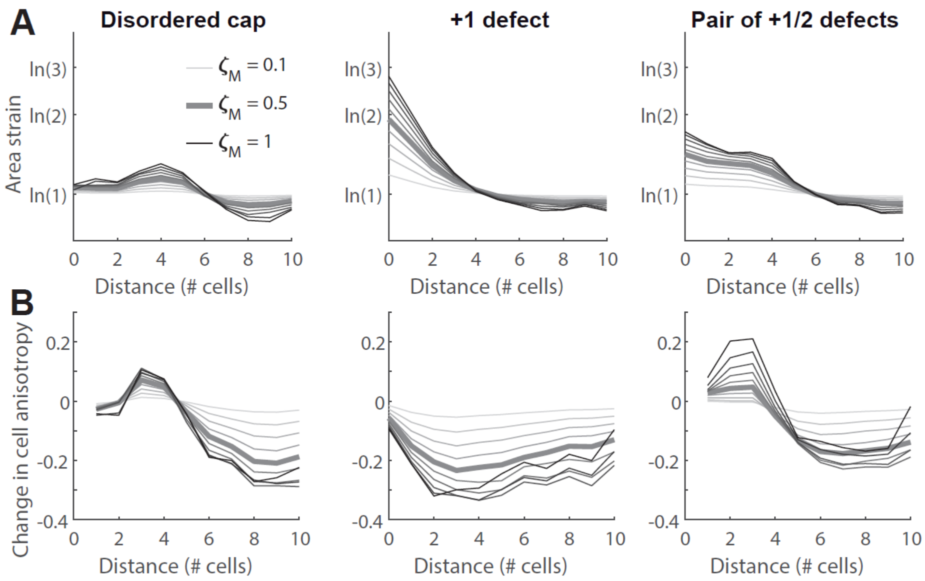


**Figure S4. Results of model simulations of mechanical strain focusing at actin foci as a function of activation amplitude.** Plots showing the average cell area strain (A) and change in cell anisotropy (B) during the peak of an activation event, as a function of distance in cells from the focus region in model simulations. The cell area strain is defined as $\epsilon_c = \ln\left(A_c/A_c^{ref}\right)$, where $A_c$ is the cell area at the peak of the event and $A_c^{ref}$ is the initial cell area. The change in cell anisotropy is defined as $\Delta Q_{rr} = Q_{rr} - Q_{rr}^{ref}$, where $Q_{rr}$ and $Q_{rr}^{ref}$ are the radial components (relative to the focus point) of the cell elongation tensor $\boldsymbol{Q}$ in the strained and relaxed states, respectively. Simulations of regenerating fragments at early stages of the regeneration process show cell stretching around the foci in the disordered cap regions (left panels). Simulations of regenerating fragments at later stages of the regeneration process show larger stretching around the +1 defect on one side (middle panels) and more moderate stretching at the pair of +½ defects at the opposite focus (right panels). The different lines depict the simulation results for different values of the activation amplitude $\bar{\zeta}_M$, going from 0.1 (light gray) to 1 (black) in steps of 0.1. Simulation results for $\bar{\zeta}_M = 0.5$, which is the value used in Figs. 5 and 7, are denoted by a thick line.

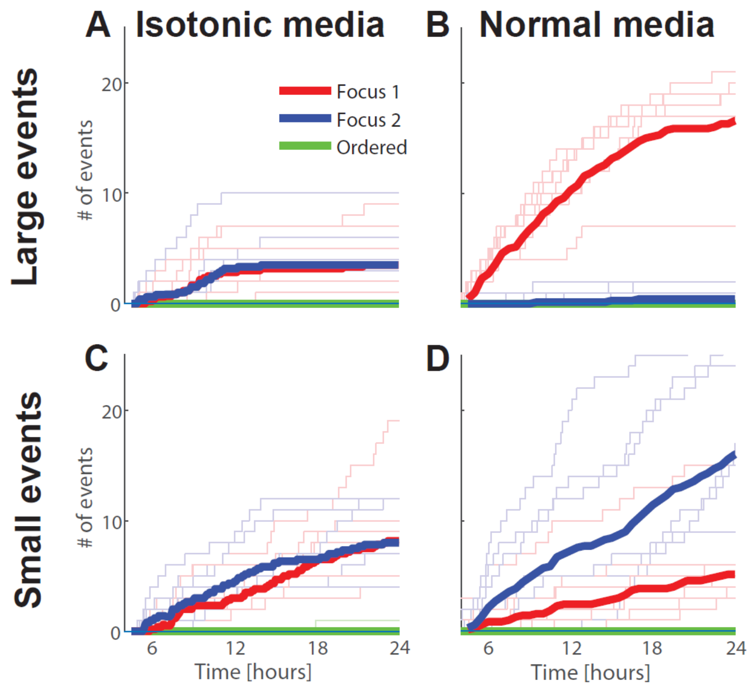


**Figure S5. Analysis of cumulative deformation events in isotonic media.** Comparison of the distribution of larger (A-B) and smaller (C-D) stretching and rupture events in isotonic media (A,C) to normal media (B,D). Events are manually characterized as in Figure 2 (Methods). Large events are defined as events in which cells stretch more than ~2-fold in area or the tissue ruptures, whereas small events are defined as events with global cell strain changes with less than a ~2-fold increase in area. The data is collected from 6 high-resolution movies of regenerating fragments in isotonic media imaged from both sides using an up-and-under microscopy setup (Methods). Events are tracked during the first 24 hours of the regeneration process, starting ~4 hours after excision (after the spheroids fold and seal). The thin lines depict results for individual movies, and the thick lines denote the mean over all samples. (A) Plot showing the cumulative number of large stretching and rupture events observed as a function of the location of the event site. (B) Plot showing the cumulative number of large stretching and rupture events as in (A) collected from 7 high-resolution up-and-under movies of regenerating Hydra in normal media (data taken from Figure 2). (C-D) Plots as in (A-B) for small stretching events.

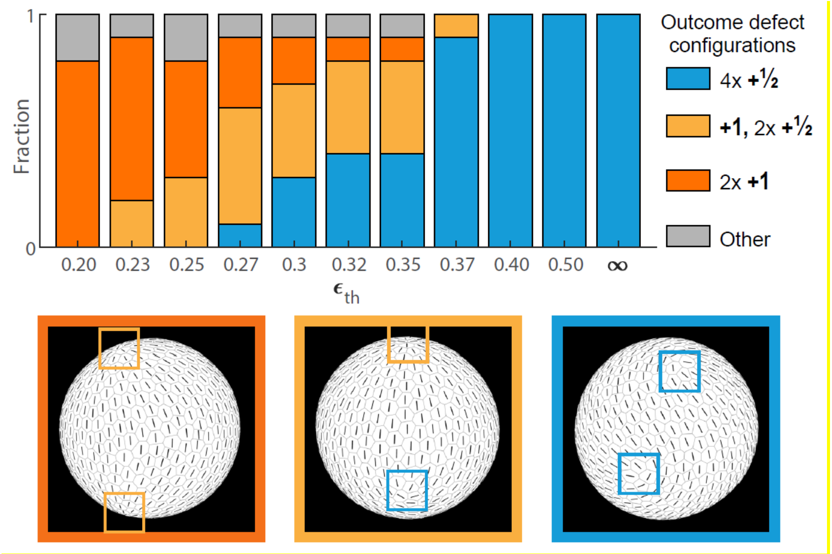


**Figure S6. Outcome defect configurations as a function of strain threshold in model simulations.** Regenerating Hydra tissues are simulated using the dynamical equations for the vertex positions, the nematic field and the morphogen concentration (Methods). The cell area strain threshold for morphogen production ($\varepsilon_{th}$) is varied from 0.2 to 0. 5 or taken to be infinite (i.e. with no morphogen production), while keeping all other model parameters fixed. The simulations are initiated with a large disordered domain of net charge +2 that is surrounded by a fully ordered region, emulating the actin fiber organization in regenerating fragments at early stages of the regeneration process (as in Fig. 5B) and no morphogen. 10 different realizations of the nematic orientations in the initial disordered region are used (see Methods). The simulation results for each value of the strain threshold for the 10 different realizations of the initial conditions, are summarized in a bar plot showing the outcome defect configurations as a function of strain threshold. Examples of the different outcome defect configurations are shown below. In the absence of morphogen ($\varepsilon_{th}$=∞) or at high strain threshold values, the nematic self-organizes into a configuration with 4 +½ defects. At lower strain threshold values, aster-shaped +1 defect are stabilized at sites of morphogen peaks.

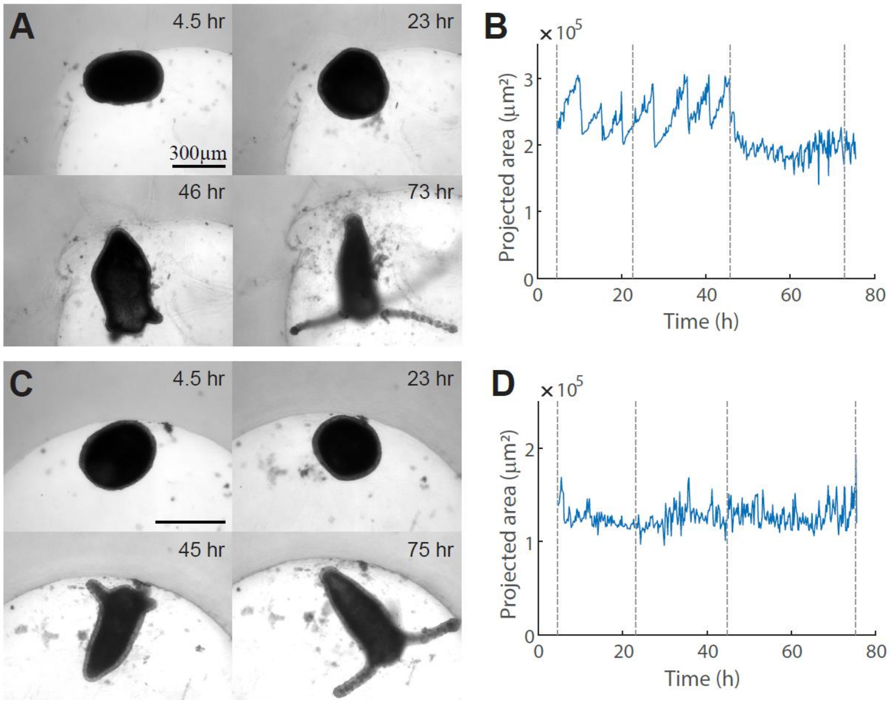


**Figure S7. Dynamics of regenerating Hydra tissue fragments.** (A-D) Examples of the overall tissue dynamics of regenerating Hydra fragments in solution. (A,C) Images from bright-field movies of regenerating Hydra tissue fragments (Methods). (B,D) Graphs of the projected area of the regenerating tissues as a function of time for the movies shown in A and C, respectively. The dashed lines indicate the time of the images shown in A and C. While most samples exhibit a characteristic saw-tooth pattern (as in B), a fraction of samples (3 out of 26 examined) do not display this pattern, yet regenerate normally (as in C,D). Similarly, we have previously shown that tissue pieces regenerating on thin wires that pierce the tissue and generate a leaky spot, exhibit tissue dynamics with a few or no saw-tooth yet are still able to regenerate (Livshits et al., 2017).

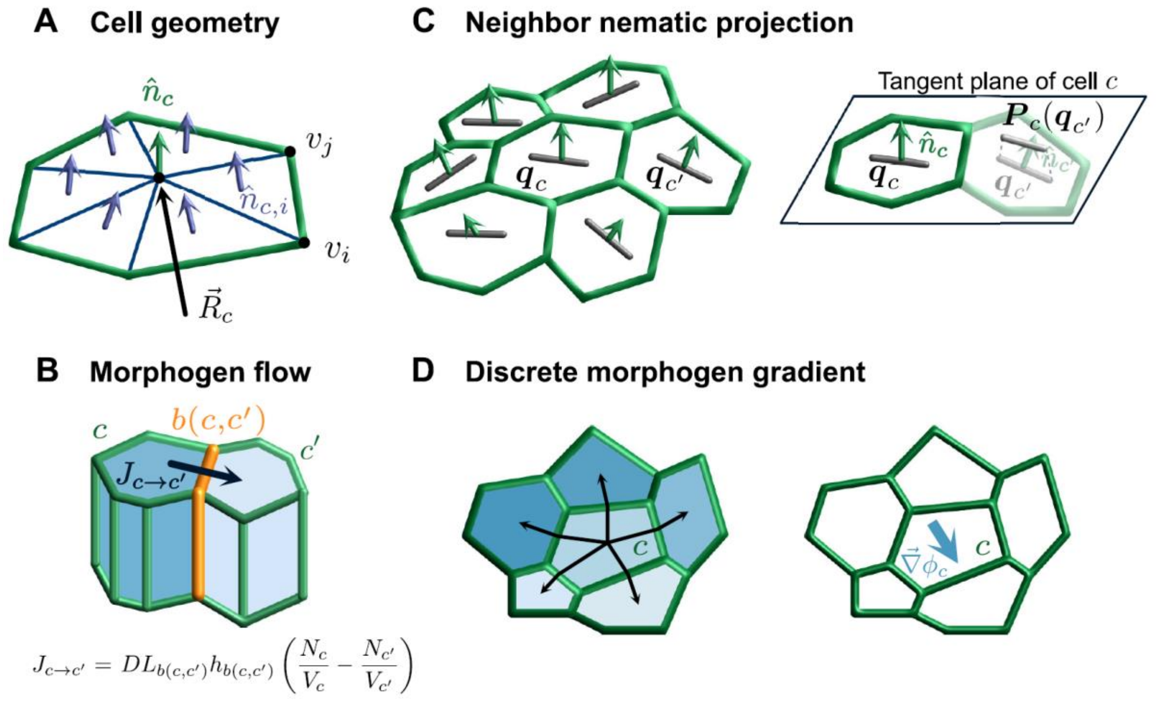


**Figure S8. Components of the mechanochemical model of Hydra regeneration.** (A) Schematics of cell triangulation and the corresponding triangle normal vectors, whose average defines the cell normal vector. (B) Illustration of the three-dimensional picture of epithelial cells. Two neighboring cells share an interface of length $L_{b(c,c')}$ and height $h_{b(c,c')}$ through which morphogen can diffuse. The diffusion flux between neighboring cells is proportional to their morphogen concentration difference and the interface area. The transport coefficient $D$ describes morphogen diffusion between the two cells. (C) Schematic of cell $c$ and its neighbors with the orientation of the nematic in each cell indicated by black bars and the cell normal vectors indicated by green arrows. In order to compute $\langle \boldsymbol{q} \rangle_c$ we calculate for the nematic tensor in each of the neighboring cells $\boldsymbol{q}_{c'}$ its projection $\boldsymbol{P}_c(\boldsymbol{q}_{c'})$ in the tangent plane of cell $c$, and then average all the projected tensor in the tangent plane. (D) Illustration of the discrete morphogen gradient in cell $c$. The morphogen concentration $\phi$ values in each cell are indicated in blue color, with darker colors corresponding to higher values. The discrete morphogen gradient in cell $c$ is defined as the vector $\vec{\nabla}\phi_{\mathrm{c}}$ that minimizes the error in the relation $\phi_{c'} = \phi_c + \vec{\nabla}\phi_{\mathrm{c}} \cdot \vec{r}_{c,c'}$ simultaneously for all neighboring cells $c'$.

### Supplementary Movie Captions

**Supplementary Movie 1: Foci of the actin fiber organization develop into topological defects localized at the regenerated head and foot regions**

Time-lapse movies of a regenerating Hydra spheroid expressing Lifeact-GFP in the ectoderm acquired simultaneously from two opposite sides of the tissue using an up-and-under spinning-disk confocal microscope (Fig. 1C; upper panels: bottom 20x objective; lower panels: top 10x objective). The two regions around the foci of the actin fiber organization were labeled by laser-induced uncaging of Abberior CAGE 552 in the folded spheroid ~3 hours after excision (Methods), allowing us to follow the two focus regions throughout the regeneration process. An aster-shaped +1 defect emerges in the focus region at the future head site (orange labels), while a pair of +½ defects emerge in the second focus at the future foot site (cyan labels). The locations of the actin foci are determined manually, and depicted by dots that are replaced by square outlines once well-defined point defects appear. The images show projected views of the ectodermal basal surface (left), the ectodermal apical surface (middle), and the basal layer with an overlay of the photoactivated dye (magenta; right). The images were centered to correct for movements of the whole tissue. The elapsed time from excision is displayed (hh:mm), and the scale bar is 100 μm.

**Supplementary Movie 2: Mechanical strain focusing at the future head region.**

High-resolution time-lapse, spinning-disk confocal movie acquired every 20 seconds of a regenerating tissue fragment expressing Lifeact-GFP in the ectoderm. The tissue spheroid is oriented with its future head region facing the objective. The images show the fiber organization at the basal surface (left) and the cellular organization at the apical surface (right). Deformation events involving mechanical strain focusing at the future head site are indicated by green frames, and are slowed down to better demonstrate the tissue deformations. The elapsed time from excision is displayed (hh:mm:ss), and the scale bar is 100 μm.

**Supplementary Movie 3: Characterization of mechanical strain around the future head site in a regenerating tissue spheroid.**

Time-lapse, spinning-disk confocal movie of a regenerating tissue fragment expressing Lifeact-GFP in the ectoderm (Fig. 3A). The tissue spheroid was initially oriented with its future head region facing the objective, showing the focus region that develops into a +1 defect (orange labels). The images show the fiber organization at the basal surface (upper left), the cellular organization at the apical surface (upper right), and cell segmentation maps depicting ectodermal cells labelled according to their area (lower left; Range (white to red): 300-1200$\mu m^2$) or cell anisotropy (lower right; Range (white to red): Q=0.1-0.65). Stretching events leading to mechanical strain focusing at the future head side are indicated by green frames, and are slowed down to better demonstrate the tissue deformations. The elapsed time from excision is displayed (hh:mm), and the scale bar is 100 μm.

**Supplementary Movie 4: Characterization of mechanical strain around the future foot region in a regenerating tissue spheroid.**

Time-lapse, spinning-disk confocal movie of a regenerating tissue fragment expressing Lifeact-GFP in the ectoderm (Fig. 3H). The tissue spheroid was oriented with its future foot region facing the objective, showing the focus region that develops into a pair of +½ defects at the future foot region (cyan labels). The images show the fiber organization at the basal surface (upper left), the cellular organization at the apical surface (upper right), and cell segmentation maps depicting ectodermal cells labelled according to their area (lower left; Range (white to red): 300-1200 $\mu m^2$) or cell anisotropy (lower right; Range (white to red): Q=0.1-0.65). The images were centered to correct for movements of the whole tissue. The elapsed time from excision is displayed (hh:mm), and the scale bar is 100 μm.

**Supplementary Movie 5: The formation of a rupture hole during a large stretching event at the focus of the actin fiber organization**

High-resolution time-lapse, spinning-disk confocal movie acquired every 20 seconds showing the formation of a rupture hole in a regenerating Hydra expressing Lifeact-GFP in the ectoderm (Fig. 4B). The rupture occurs during a large stretching event, initiating as an elongated crack along ectodermal cell-cell junctions, rapidly becoming circular, and sealing back within minutes. The movie shows additional large stretching events (indicated by green frames), with mechanical strain focusing recurring at the same location, at the focus site of the ectodermal actin fibers. The events are slowed down to better demonstrate the tissue deformations. The images show the fiber organization at the basal ectodermal surface (left) and the cellular organization at the apical ectodermal surface (right). The elapsed time from excision is displayed (hh:mm:ss), and the scale bar is 100 μm.

**Supplementary Movie 6: An extremely large rupture hole in a regenerating tissue spheroid**

Time-lapse, spinning-disk confocal movie showing a regenerating tissue spheroid expressing Lifeact-GFP in the ectoderm that experiences two extremely large rupture holes lasting for an extended period of time (~1 hour; Fig. 4C). The movie shows that the tissue spheroid is nevertheless able to successfully regenerate into a mature Hydra. The rupture holes form at the +1 defect region at the future head site (orange labels). Large deformation events involving mechanical strain focusing and rupture hole formation recur at the same location and are indicated by green frames, and are slowed down to better demonstrate the tissue deformations. Later large deformation events that are observed from the side (since the tissue rotated and the foci regions are not visible) are indicated by light green frames. The images show the fiber organization at the basal ectodermal surface (left) and the cellular organization at the apical ectodermal surface (right). The images were centered to correct for movements of the whole tissue. The elapsed time from excision is displayed (hh:mm), and the scale bar is 100 μm.

**Supplementary Movie 7: Simulation of Hydra tissue mechanics during global fiber contraction in a spheroid with a large disordered domain.**

The movie depicts the results of simulations of a transient activation of global fiber contraction in a Hydra tissue spheroid with a nematic configuration containing a large disordered domain of net charge +2 surrounded by a fully ordered region, emulating the nematic pattern at the early stages of regeneration (Fig. 5B). The tissue spheroid is modeled as a deformable shell that contains an embedded nematic field that experiences a time-dependent global activation (bottom inset). The images show cell segmentation maps depicting cells labelled according to the cell area strain defined as $\epsilon_c = \ln\left(A_c/A_c^{ref}\right)$, where $A_c$ is the cell area and $A_c^{ref}$ is the initial cell area, overlaid with lines indicating the orientation of the embedded nematic field in each cell (black- ordered region; gray- disordered regions). The spheroid is shown from a top view centered on one of the actin foci and a side view. The active stress generated within each cell is aligned with the nematic field in that cell, and global activation leads to mechanical strain focusing around the two foci.

**Supplementary Movie 8: Simulation of Hydra tissue mechanics during global fiber contraction in a spheroid with a +1 defect and a pair of +½ defects.**

The movie depicts the results of simulations of a transient activation of global fiber contraction in a Hydra tissue spheroid with a nematic configuration containing a +1 defect on one side of the spheroid and a pair of +½ defects at the opposite end, emulating the actin fiber organization in regenerating fragments at later stages of the regeneration process (Fig. 5C). The tissue spheroid is modeled as a deformable shell that contains an embedded nematic field that experiences a time-dependent global activation (bottom graph). The images show cell segmentation maps depicting cells labelled according to the cell area strain defined as $\epsilon_c = \ln\left(A_c/A_c^{ref}\right)$, where $A_c$ is the cell area and $A_c^{ref}$ is the initial cell area, overlaid with lines indicating the orientation of the embedded nematic field in each cell. The spheroid is shown from a top view centered on the aster-shaped +1 defect, and a bottom view showing the pair of +½ defects. The active stress generated within each cell is aligned with the nematic field in that cell, and global activation leads to efficient mechanical strain focusing at the +1 defect site (right), and to a lesser extent at the pair of +½ defects (left).

**Supplementary Movie 9: The effect of iCRT14 treatment on actomyosin fiber organization**.

Time-lapse, spinning-disk confocal movie of a regenerating spheroid expressing Lifeact-GFP in the ectoderm, that was treated with 5 µM iCRT14 (Fig. 6B; Methods). The tissue spheroid was initially oriented with its closure region facing the objective. The actomyosin fibers do not reform in the closure region and point defects do not emerge, whereas the inherited aligned fibers visible around the closure region are maintained. The images show the fiber organization at the basal ectodermal surface (left) and the cellular organization at the apical ectodermal surface (right). The images were centered to correct for movements of the whole tissue. The elapsed time from excision is displayed (hh:mm), and the scale bar is 100 μm.

**Supplementary Movie 10: The effect of elevated medium osmolarity on actin fiber organization.**

Time-lapse, spinning-disk confocal movie of a regenerating tissue spheroid expressing Lifeact-GFP in the ectoderm, placed in an isotonic medium containing 70 mM sucrose (Fig. 6E; Methods). The movie shows a disordered region that rapidly orders and develops a +1 defect (orange labels), that subsequently unbinds into a pair of +½ defects (cyan labels). The tissue spheroid stabilizes in a 4x +½ defect configuration and elongates along the fibers' direction but does not regenerate. The images show the fiber organization at the basal ectodermal surface (left) and the cellular organization at the apical ectodermal surface (right). The images were centered to correct for movements of the whole tissue. The elapsed time from excision is displayed (hh:mm), and the scale bar is 100 μm.

**Supplementary Movie 11: Simulation of Hydra tissue regeneration resulting in an asymmetric defect configuration with a +1 defect and a pair of +½ defects.**

Simulation results showing a regenerating tissue spheroid initiated with a partially ordered fiber configuration and no morphogen (Fig. 7C; see SI for details). The tissue deformations and the dynamics of the nematic and morphogen concentration fields are modeled through 30 recurring global fiber contraction events (the cycle is indicated). The spheroid is shown from a top view centered on one of the actin foci and a side view. The tissue self-organizes into a configuration with an aster-shaped +1 defect colocalized with a morphogen peak at one end of the spheroid (emulating the organizer emerging at the future head site), and a pair of +½ defects at the opposite side.

**Supplementary Movie 12- Simulation of Hydra tissue regeneration resulting in a symmetric defect configuration with two +1 defects.**

Simulation results showing a regenerating tissue spheroid initiated with a partially ordered fiber configuration and no morphogen (Fig. 7C see SI for details). The tissue deformations and the dynamics of the nematic and morphogen concentration fields are modeled through 30 recurring global fiber contraction events (the cycle is indicated). The spheroid is shown from a top view centered on one of the actin foci and a side view. The tissue self-organizes into a configuration with two-aster shaped +1 defects colocalized with two morphogen peaks.

**Supplementary Movie 13: Simulation of Hydra tissue regeneration without morphogen.**

Simulation results without coupling to a morphogen field showing a regenerating tissue spheroid initiated with a partially ordered fiber configuration (Fig. 7C; see SI for details). The tissue deformations and the dynamics of the nematic are modeled through 30 recurring global fiber contraction events (the cycle is indicated). The spheroid is shown from a top view centered on one of the actin foci and a side view. The tissue self-organizes into a configuration with four +½ defects.
